# Endocardial TIE1 synergizes with TIE2 to regulate the atrial internal muscular network assembly

**DOI:** 10.64898/2026.02.26.708190

**Authors:** Kai Ding, Beibei Xu, Xinhao Yu, Xiwen Jia, Taotao Li, Xin Shen, Junda Li, Xudong Cao, Yahui Liu, Zhen Zhang, Yulong He

**Affiliations:** Cyrus Tang Hematology Center, Collaborative Innovation Center of Hematology, National Clinical Research Center for Hematologic Diseases, State Key Laboratory of Radiation Medicine and Protection, Cam-Su Genomic Resources Center, Suzhou Medical College of Soochow University, Suzhou 215123, China; Pediatric Translational Medicine Institute and Pediatric Congenital Heart Disease Institute, Shanghai Children’s Medical Center, Shanghai Jiao Tong University School of Medicine, Shanghai 200127, China; Shanghai Collaborative Innovative Center of Intelligent Medical Device and Active Health, Shanghai University of Medicine & Health Sciences, Shanghai 201318, China

**Keywords:** TIE1, TIE2, endocardium, trabeculation, atrial internal muscular network, chamber morphogenesis

## Abstract

Atrial cardiomyopathy is characterized by altered atrial structures, yet its underlying genetic basis remains inadequately explored. TIE1 mutations are reported in lymphedema patients, but whether they exhibit cardiac defects alongside lymphatic abnormalities remains unknown. Here, single-cell RNA sequencing showed high *Tie1* and *Tek* expression in atrial endocardial cells. TIE1 deficiency severely disrupted atrial morphogenesis, with only minor ventricular defects. Bulk RNA-seq showed reduced transcripts for endothelial development and cardiac trabeculation in *Tie1* mutants. Separate RNA-seq analysis of the atria and ventricles further confirmed these findings, revealing a more pronounced downregulation of trabecular genes, including *Tek*, in the atria upon *Tie1* loss. Endothelial deletion of *Tek* impaired trabeculation, particularly in atria. Consistently, *Tie1* loss combined with a single null *Tek* allele disrupted both atrial and ventricular trabeculation. Strikingly, atrial morphogenesis defects appeared within 48 hours of induced endothelial *Tie1* loss coupled with *Tek* heterozygous deletion, implying a critical endocardial role in organizing the atrial muscular network. The synergy of TIE1 and TIE2 in remodeling atrial trabeculae was confirmed postnatally, whereas TIE1 insufficiency alone had no obvious effect. Together, TIE1 is differentially required for atrial versus ventricular development, acting synergistically with TIE2 to regulate endocardial cell-coordinated assembly of atrial internal muscular network.

## Introduction

The endocardium, as a specialized type of endothelial cells (EC) lining the cardiac lumen, plays an important role in vertebrate heart morphogenesis [1, 2]. Cardiac trabeculation relies on the coordinated interaction of endocardium and myocardium [3, 4]. The endocardium senses mechanical forces and provides signals for the myocardial morphogenesis [5]. When endocardial cells fail to form buds, the assembly and extension process of trabeculae are disrupted [1, 6]. Defective trabecular formation could lead to embryonic lethality or congenital heart disease [7–9]. Cell lineage studies have also shown that the endocardium is an important source of the coronary vascular endothelial cells, and gives rise to most mesenchymal cells that constitute the cardiac cushions and subsequently contribute to the heart valves [2]. Atrial internal muscular networks, mainly pectinate muscles running in an anterolateral direction, are found on the anterior surface of both right and left atrial walls and corresponding auricles. In contrast to the better understanding of ventricular trabecula morphogenesis, the dynamic process and regulatory networks of atrial trabeculation are inadequately elucidated. Likewise, the atrial cardiomyopathy is characterized by altered atrial structure as well as function, and pathological mechanisms underlying the disorders are largely unknown.

Cardiac morphogenesis is a complex process involving multiple pathways to orchestrate the reciprocal paracrine signaling between the endocardial and other cardiac cells. Specifically, myocardial VEGFA activates VEGFR2 signaling for coronary vascular growth and also participates in the regulation of cardiac trabeculation [10]. Endocardial cell-derived NRG1 regulates myocardium development via its receptors ERBB2/4 in cardiomyocytes. Mice null for *Nrg1* or its receptors display hypoplastic ventricular wall lacking normal trabeculation [11]. The aberrant NOTCH1 signaling was found to disrupt the heart morphogenesis [12, 13]. NRG1 has been shown to regulate VEGFA expression in the myocardium, and VEGFA-VEGFR2 signaling is required for the NOTCH1 pathway restriction in endocardium during trabeculation. NOTCH1 signaling was shown to promote extracellular matrix (ECM) degradation during the formation of endocardial projections while NRG1 promoted myocardial ECM synthesis for trabecular rearrangement during the process of cardiac trabeculation [14]. It has recently been shown that endocardium and myocardium interacted directly through tunneling nanotube–like structures extending from cardiomyocytes to endocardial cells, contributing to NOTCH1 activation [15]. Factors involved in the synthesis or degradation of cardiac jelly, a layer of extracellular matrix between endocardium and myocardium, also play essential roles in the process of chamber morphogenesis, including glycosaminoglycan proteins such as versican and ADAMTS family proteins [16, 17]. The ANGPT1-TIE2 pathway plays a crucial role in cardiovascular development [18–21]. ANGPT1 is expressed by cardiomyocytes, particularly in atrial myocardium and ventricular trabeculae, and participates in the atrial morphogenesis by regulating the spatiotemporal degradation of cardiac jelly [18]. The endocardial specific deletion of *Tek* resulted in the fewer but thicker ventricular trabeculae as well as impaired endocardial sprouting, potentially via the paracrine suppression of retinoic acid signaling and proliferation in trabecular cardiomyocytes [21].

Endothelial TIE1 shares high homology with TIE2 and could form TIE1/TIE2 heterodimers [22]. TIE1 has been shown to participate in the regulation of blood vascular and lymphatic network formation and also aortic valve development [23–27]. It has recently been shown that TIE1 and TIE2, via the PI3K-AKT-mediated signaling pathway for the protein stability of COUP-TFII, regulate venous specification [25, 28]. Loss of the transcription factor COUP-TFII was reported to disrupt both the vein and atrial development [29–31]. *Tie1* mutations or variants have also been linked to lymphatic disorders [32, 33]. It remains to be characterized whether Tie1 mutations could affect heart structure and function, and how TIE1 participates in the process of cardiac wall morphogenesis. To investigate the role of TIE1 and TIE2 in the process of endocardial cell-mediated chamber formation, we employed genetic mouse models targeting *Tie1*, *Tek* (encoding TIE2) or both at different developmental stages. Based on the analysis of the published single cell RNA sequencing data (scRNA-seq) by Feng et al. [34] and the RNA-seq analysis of atria and ventricles from the *Tie1* mutant and littermate control mice, we found in this study that *Tie1* and *Tek* expression were higher in atrial than ventricular endocardial cells. TIE1 is differentially required for the atrial and ventricular trabeculation, acting in synergy with TIE2.

## Materials and Methods

### Mouse models

All animal procedures performed conform the guidelines from Directive 2010/63/EU of the European Parliament on the protection of animals used for scientific purposes; and the experiment protocols were approved by the Animal Care Committee of Soochow and Nanjing University Animal Center (MARC-AP#YH2 /SUDA20250507A03). All the mice used in this study were housed in a SPF (specific pathogen free) animal facility with a 12/12 hours dark/light cycle and were free to food and water access. Normal mouse diet (Suzhou Shuangshi Experimental Animal Feed Technology, Co, Ltd) and cage bedding (Suzhou Baitai Laboratory Equipment, Co, Ltd) were used. Two genetically modified mouse models targeting *Tie1* gene were employed in this study. One mouse line targets TIE1 intracellular kinase domain (ICD), with exon 15 and exon 16 floxed, *Tie1ICD^Flox/Flox^*. The other line is a knockout first mouse model established from EUCOMM embryonic stem cells (EPD0735-3B07) targeting *Tie1* gene (*Tie1^tm1a/tm1a^*), in which targeting cassette is recombined downstream of exon 7 (with exon 8 floxed) [25, 27]. *Tek* knockout mouse model was generated as previously reported [28] and was crossbred with *Tie1*^Δ*ICD/*Δ*ICD*^ mouse model to obtain *Tie1* and *Tek* double knockout mice (*Tie1*^Δ*ICD/*Δ*ICD*^*;Tek^+/-^*). To generate mice with endothelial cell–specific gene deletion mouse models, we employed the *Cdh5-Cre^ERT2^* mouse line [35]. In all the phenotype analysis, wildtype or heterozygous littermates were used as controls. For the genotyping of *Tie1* and *Tek* mouse lines, the primers used were as previously described [25, 27]. The genetic background of *Tie1^tm1a/tm1a^* is on C57BL/6 N, and the other lines are on C57BL/6J or C57BL/6J/SV129. Mice were sacrificed by asphyxiation with rising concentration of carbon dioxide gas, followed by cervical dislocation and tissue collection.

### Induced gene deletion

Induction of gene deletion was performed as previously described by the tamoxifen treatment [25, 36]. Briefly, pregnant mice were treated with tamoxifen (Sigma-Aldrich, T5648-5G) at E10.5-E11.5, E12.5-14.5 or E12.5-15.5 (1 mg/per mouse for 2-4 consecutive days by intraperitoneal injection) and embryos were collected for analysis at the specified stages. For the postnatal studies, new-born pups were treated by four daily intragastric injections of tamoxifen starting from postnatal day 1 (P1, 60 μg/per mice daily). Tissues were collected for analysis at postnatal day 21. The genotypes of the *Tie1/Tek* knockout and control mice are as follows: 1) *Tie1ICD^Flox/-^;Cdh5-Cre^ERT2^*named as *Tie1ICD^iECKO^*, littermates *Tie1ICD^Flox/+^*or *Tie1ICD^Flox/+^;Cdh5-Cre^ERT2^* mice named as *Control*. 2) *Tek^Flox/-^;Cdh5-Cre^ERT2^* named as *Tek^iECKO^*, littermates *Tek^Flox/+^* or *Tek^Flox/+^;Cdh5-Cre^ERT2^* mice named as *Control*. 3) *Tie1ICD^Flox/-^;Cdh5-Cre^ERT2^;Tek^+/-^*named as *Tie1ICD^iECKO^;Tek^+/-^*, littermates *Tie1ICD^Flox/+^;Cdh5-Cre^ERT2^;Tek^+/-^*, *Tie1ICD^Flox/+^;Tek^+/-^*, *Tie1ICD^Flox/+^;Cdh5-Cre^ERT2^;Tek^+/+^*or *Tie1ICD^Flox/+^;Tek^+/+^* named as *Control*. In this study, the *Tie1/Tek* double mutant mice refer to *Tie1*^Δ*ICD/*Δ*ICD*^*;Tek^+/-^* or *Tie1ICD^iECKO^;Tek^+/-^*, which harbor a *Tie1* null mutation or an endothelial-specific deletion alongside the heterozygous knockout of *Tek*.

### Embryo preparation

To investigate the developmental process of cardiac chamber formation, female mice were mated in the late afternoon and vaginal plugs were checked in the morning of the following day. The embryonic stages are estimated considering midday of the day on which the vaginal plug is present as embryonic day 0.5 (E0.5). Tail tips from mutant and control embryos were used for genotyping, and heart tissues from the embryos were collected for the subsequent histology and RNA analysis. For the RNA preparation, heart tissues were harvested and placed in cold PBS. The residual blood from the hearts was cleaned and heart tissues were snap-frozen in liquid nitrogen. For the separate analysis of atria and ventricles, heart tissues were collected and dissected before snap-freezing.

### Immunostaining

For the frozen tissue section staining, tissues were collected and fixed in 4% paraformaldehyde for 2 hours at 4°C, followed by the incubation in 20% sucrose overnight before being embedded in optimal cutting temperature (OCT) compound. Frozen sections (10 µm) were used for the immunostaining analysis of cardiac chamber structures and coronary vessels. For the visualization of atrial internal muscular network-trabeculae structures, the whole-mount immunostaining was performed. Heart tissues were fixed in 4% paraformaldehyde overnight at 4°C, followed by blocking with 3%(w/v) skim milk in PBS-TX (0.3% Triton X-100) and incubating with the primary antibodies. The antibodies used were: hamster anti-mouse PECAM1(MAB1398Z; Millipore); rat anti-mouse CD31 (553370; BD Pharmigen), rat anti-mouse Endomucin (14-5851; eBioscience), goat-anti-mouse DLL4 (AF1389; R&D) and goat anti-human TIE1 (AF619; R&D). Appropriate Alexa 488(Invitrogen), Cy3(Jackson ImmunoResearch) conjugated secondary antibodies were used. All fluorescently labeled samples were mounted and analyzed with a confocal microscope (Olympus Fluoview 3000) or Leica MZ16F fluorescent dissection microscope. For comparison, the parameters were kept consistent for all the confocal microscopic imaging in this study.

### Bulk RNA-seq analysis of atria and ventricles

For the bulk RNA-seq analysis of heart tissue, the whole hearts from mutant and control embryos were collected. Atria and ventricles were separated and tissues from two embryos with the same genotype (*Tie1^tm1a/tm1a^*and *control*) were pooled for the RNA preparation and further analysis. According to the manufacturer’s instructions, total RNA was extracted from the tissues using Trizol (Invitrogen). The procedures for the RNA-seq analysis were performed as previously reported [25]. Briefly, sequencing was performed on DNBSEQ platform with PE150 (read length, BGI-Shenzhen, China), and the sequencing data were processed according to the predefined analysis pipeline. The differential expression analysis was conducted using the R package DESeq2 (v 1.42.1). Functional enrichment analysis was performed using clusterProfiler (v4.10.1) and org.Mm.eg.db (v 3.18.0). The hallmark gene sets used in the GSEA (Gene Set Enrichment Analysis), including the hallmarks for trabeculation and EC migration, are mainly defined according to the transcriptome atlas of murine endothelial cells, the molecular signatures database (mh.all.v2024.1.Mm.symbols) and the related GO terms [25, 37, 38]. The raw and processed RNA-seq data will be deposited in the GEO database after the publication.

### Analysis of embryonic cardiac single-cell RNA-seq data

The single-cell RNA sequencing data (GSE193346) by Feng et al. [34] were processed and analyzed with the Seurat package (v5.2.1) following the same parameters and criteria as described to ensure consistency and comparability. Major cell types were annotated based on established marker genes: pan-cardiomyocyte (*Ttn*), atrial cardiomyocytes (*Sln*), ventricular cardiomyocyte (*Myl2*), pan-endothelial cells (*Pecam1*), endocardial cells (*Npr3*), coronary endothelial cells (*Fabp4*), epicardial cells (*Wt1*, *Tbx18*, and *Aldh1a2*), fibroblast-like cells (*Postn*), mural cells (*Pdgfrb*), red blood cells (*Hba-a1*), macrophages (*Adgre1*). Following the identification of endocardial cells, subsequent analysis was performed using the Seurat package and normalized by the “LogNormalize” method, highly variable genes were identified with the ‘FindVariableFeatures” function (selection.method = “vst”, nfeatures = 2000). The optimal number of principal components (dims = 1:22) was determined by the ElbowPlot method. Cells were then clustered using the “FindNeighbors” and “FindClusters” functions (dims= 1:16, resolution = 0.5). Finally, Unsupervised clustering of cells will be performed by “RunUMAP” and gene expression was visualized with “FeaturePlot”. Ligand–receptor interactions between cardiomyocytes and endocardial cells during the embryonic stage (E10.5-E18.5) were identified using the CellChat package (v1.6.1). Atrial and ventricular cell subsets were analyzed separately using the mouse ligand-receptor database (CellChatDB.mouse) and communication probabilities were computed (min.cells = 10) to identify significant interactions.

### Quantitative analysis of cardiac wall and trabeculae

For the quantification of trabecular structures in atria and ventricles, the EMCN positive areas was measured. The PECAM1 positive area was quantified to indicate the coronary vessels within the ventricular wall. The cardiac wall thickness was also measured as described in the **Figure 1–figure supplement** 2. For the quantitative analysis of dorsal coronary veins, images were taken under a fluorescence microscope and kept constant near the venous sinus for all of the samples. Horizontal lines were evenly laid on the images, and the diameters of veins crossed with these lines were measured and analyzed using Image Pro Plus (Media Cybernetics, Inc., Bethesda, MD). The ratio of heart to embryo length was used to represent heart size. The starting point for the heart measurement was the junction of aorta and pulmonary artery, and the junction of left and right ventricles at the apex as the end point. All the quantification was measured and analyzed by using Image Pro Plus. Furthermore, we also performed the measurement of cardiac trabecular complexity by the Fractal analysis. Fractal dimensions were derived from confocal images of hearts from *Tie1* mutants and control mice. Quantification was carried out using ImageJ with the Fraclac plugin (including 6 grid orientations).

### Statistical analysis

Data analyses were performed using GraphPad Prism (version 8.0). For the 2-group comparison, the unpaired t test was performed with Welch correction if data passed the D’Agostino-Pearson normality test, or the unpaired nonparametric Mann-Whitney U test was applied. Data are expressed as mean ± SD. All statistical tests were 2-sided.

## Results

### Enriched expression of *Tie1* and *Tek* in atrial compared with ventricular endocardial cells

To dissect the transcriptional dynamics of key endothelial regulators during the murine heart development, we analyzed the published single-cell transcriptomic dataset (GSE193346) spanning the murine embryonic development (E10.5-E18.5) [34]. We found that *Tie1*, and *Tek* were highly expressed in endocardial cells (*Pecam1*^+^, *Npr3^+^*) while the coronary EC gene such as the coronary endothelial marker *Fabp4* was lowly expressed in endocardial cells (**Figure 1A-B, Figure 1– figure supplement** 1A-C). Interestingly, we found that the transcript levels of *Tie1*, and *Tek* were higher in the atrial endocardial cells than those of ventricles, especially at embryonic stages up to E14.5 when the structures of four-chambered hearts were established (Table 1). In contrast, it appears that the *Angpt1* transcript level was higher in atrial than ventricular cardiomyocytes (**Figure 1A-B**, Table 1).

**Figure 1.**
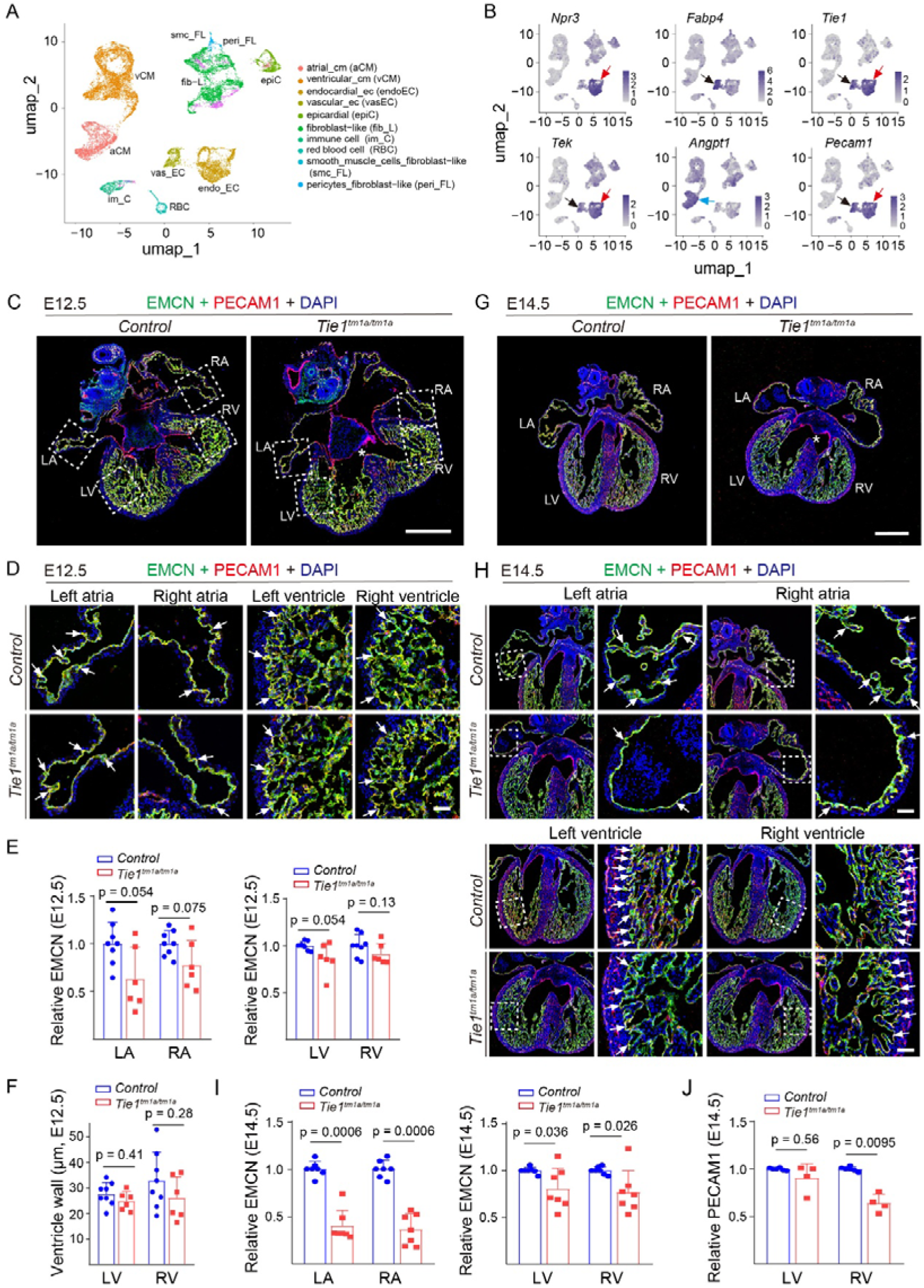
Single-cell RNA-seq analysis of *Tie1/Tek* and its ligand *Angpt1* expression in endocardial cells and its role in embryonic cardiac trabeculation. **A-B.** Analysis of the single-cell RNA-seq data of embryonic hearts from E10.5 to E18.5 by Feng et al. Nat Commun 2022 (DOI: 10.1038/s41467-022-35691-7). UMAP visualization of 10 distinct cell clusters in the murine hearts during embryogenesis (E10.5 to E18.5, A). *Tie1* and *Tek* are highly expressed in the endocardial cell population (*Npr3*⁺) as shown in feature plots. *Angpt1* is highly expressed in the atrial cardiomyocytes (B). Black arrows point to the coronary EC population (*Fabp4*^+^). Red arrows point to the endocardial cell population (*Npr3*⁺). Blue arrows point to the atrial cardiomyocytes. The quantitative results are shown in Table 1. **C-J**. Analysis of heart trabeculae of *Tie1^tm1a/tm1a^* and littermate controls by immunostaining for EMCN (endomucin; green), PECAM1 (platelet endothelial cell adhesion molecule 1; red) and DAPI (blue) at E12.5 (**C-D**) and E14.5 (**G-H**). Note that *Tie1^tm1a/tm1a^* mice displayed defective trabeculation in the atrium (white arrows) and sparse trabecular structures in the ventricles (white arrows) at E12.5, with more severe defects by E14.5, lacking atrial trabeculae (white arrows) and sparser ventricular trabeculae (white arrows). Asterisks in C and G point to the delayed formation of interventricular septum in *Tie1^tm1a/tm1a^* mice. Arrows point to the trabecula-associated endocardium. **E-F.** Quantification of the EMCN-positive area in the atrial and ventricular trabeculae at E12.5 (**E**, LA: *Tie1^tm1a/tm1a^*: 0.63±0.33, n=6; *Control*: 1.00±0.22, n=8; *P*=0.054. RA: *Tie1^tm1a/tm1a^*: 0.77±0.26, n=6; *Control*: 1.00±0.13, n=8; *P*=0.075. LV: *Tie1^tm1a/tm1a^*: 0.86±0.16, n=6; *Control*: 1.00±0.046, n=8; *P*=0.054. RV: *Tie1^tm1a/tm1a^*: 0.91±0.11, n=6; *Control*: 1.00±0.12, n=8; *P*=0.13.). Quantification of the ventricular wall thickness (**F**, LV: *Tie1^tm1a/tm1a^*: 24.87±3.70 µm, n=6; *Control*: 27.67±4.52 µm, n=8; *P*=0.41. RV: *Tie1^tm1a/tm1a^*: 26.01±8.42 µm, n=6; *Control*: 32.84±11.21 µm, n=8; *P*=0.28.). **I-J**. Quantification of the EMCN-positive area in the atrial and ventricular trabeculae at E14.5 (**I**, LA: *Tie1^tm1a/tm1a^*: 0.40±0.16, n=7; *Control*: 1.00±0.079, n=7; *P*=0.00060. RA: *Tie1^tm1a/tm1a^*: 0.36±0.17, n=7; *Control*: 1.00±0.092, n=7; *P*=0.00060. LV: *Tie1^tm1a/tm1a^*: 0.81±0.22, n=7; *Control*: 1.00±0.033, n=7; *P*=0.036. RV: *Tie1^tm1a/tm1a^*: 0.77±0.23, n=7; *Control*: 1.00±0.040, n=7; *P*=0.026.). Quantification of the PECAM1-positive area in ventricular walls (**J**, LV: *Tie1^tm1a/tm1a^*: 0.91±0.15, n=4; *Control*: 1.00±0.013, n=6; *P*=0.56. RV: *Tie1^tm1a/tm1a^*: 0.64±0.096, n=4; *Control*: 1.00±0.024, n=6; *P*=0.0095.). The quantification data of mutant mice was normalized against that of littermate control mice. In E-J, statistical analysis was performed using the unpaired nonparametric Mann-Whitney U test. Values are represented as means ± SD of at least three independent technical replicates. LA:Left atria, RA:Right atria, LV:Left ventricle, RV: Right ventricle; Scale bar: 400 μm in C and G, 50 μm in D and H.

**Table 1.** The transcript levels of *Tie1*, *Tek* in atrial and ventricular endocardial cells and *Angpt1* in atrial and ventricular cardiomyocytes during the embryonic stage from E10.5-E18.5 (based on the published scRNA-seq data, Feng et al., Nat Commun 2022; DOI: 10.1038/s41467-022-35691-7). Values are represented as means ± SD and the number referred to cell counts.

|  | *Tie1* |  | *Tek* |  | *Angpt1* |  |
| --- | --- | --- | --- | --- | --- | --- |
|  | Atria<br>(mean±sd) | Ventricle<br>(mean±sd) | Atria<br>(mean±sd) | Ventricle<br>(mean±sd) | Atria<br>(mean±sd) | Ventricle<br>(mean±sd) |
| E10.5 | 1.34±0.50<br>(n = 9) | 1.17±0.39<br>(n = 16) | 1.37±0.54<br>(n = 10) | 1.31±0.50<br>(n = 17) | 0.53±0.49<br>(n = 50) | 0.28±0.41<br>(n = 197) |
| E12.5 | 1.54±0.43<br>(n = 5) | 1.27±0.47<br>(n = 220) | 1.26±0.50<br>(n = 4) | 1.17±0.48<br>(n = 215) | 0.95±1.19<br>(n = 6) | 0.29±0.40<br>(n = 665) |
| E14.5 | 1.60±0.49<br>(n = 9) | 1.41±0.48<br>(n = 72) | 1.43±0.39<br>(n = 8) | 1.37±0.48<br>(n = 61) | 1.13±0.45<br>(n = 47) | 0.18±0.34<br>(n = 322) |
| E16.5 | 1.41±0.47<br>(n = 89) | 1.35±0.49<br>(n = 35) | 1.18±0.45<br>(n = 84) | 1.49±0.45<br>(n = 33) | 1.41±0.55<br>(n = 404) | 0.17±0.41<br>(n = 112) |
| E18.5 | 1.26±0.40<br>(n = 137) | 1.35±0.47<br>(n = 30) | 1.15±0.45<br>(n = 136) | 1.24±0.43<br>(n = 25) | 1.15±0.67<br>(n = 455) | 0.098±0.28<br>(n = 277) |

To investigate mechanisms underlying the murine atrial trabeculation, we first analyzed the dynamic processes of cardiac chamber morphogenesis by the immunostaining analysis of endomucin (EMCN) for trabeculae and PECAM1 for coronary vessels (embryonic day 10.5-E18.5). The quantification methods for atrial and ventricular trabecular areas, wall thickness and vessel density were shown in **Figure 1–figure supplement** 2A. As shown in **Figure 1–figure supplement** 2B, the heart tube became looped to form the preliminary chamber structures at E10.5, including the initiation of interventricular septum. While there appeared to have an undulating, wave-like contour or the budding of endocardium in atria (E10.5), trabecular structures were readily detectable in ventricles at this stage. Therefore, atrial wall morphogenesis occurs later than that of ventricles, which initiates the trabeculation process around E8.5 when endocardial cells migrate through cardiac jelly to form anchor points with compact myocardium [14]. We found that pectinate muscles, defined as the atrial trabeculae, became detectable in the atria at E12.5 (**Figure 1–figure supplement** 2B). The formation of interventricular septum was completed by E14.5, establishing the four-chambered heart. By E16.5 or later stages (**Figure 1–figure supplement** 2C), the atrial and ventricular trabeculae gradually formed a complex network. As shown in **Figure 1–figure supplement** 2D-G (Supplement Table 1), the trabecular areas in both the atria and ventricles gradually increased, being greater in the right atria (RA) than that in the left atria (LA, **Figure 1–figure supplement** 2D). There was no obvious difference in the trabecular areas between the left and right ventricles (**Figure 1–figure supplement** 2E). The ventricular wall thickness increased rapidly after E12.5, with the left ventricular wall being thicker than that of the right ventricular wall by E18.5 (**Figure 1–figure supplement** 2F), accompanied by the increase of coronary vessel density (**Figure 1–figure supplement** 2G).

### Differential requirement of TIE1 in atrial versus ventricular trabeculation

To study the role and mechanism underlying TIE1 function in cardiac chamber morphogenesis, we employed two genetically modified mouse models targeting *Tie1*, including *Tie1^tm1a/tm1a^* and *Tie1*^Δ*ICD/*Δ*ICD*^ [25, 27]. The effects of *Tie1* gene deletion on atrial and ventricular trabeculae were analyzed at different developmental stages (including E12.5, E14.5 and E17.5). TIE1 deficiency (*Tie1^tm1a/tm1a^*) led to an impaired trabeculation in the atria, characterized by a decrease of trabecular structures at E12.5, while the formation of ventricular trabeculae was not obviously affected at the same stage (**Figure 1C-D**). Shown in **Figure 1E** was the quantification of trabecular areas in atria and ventricles. The thickness of left and right ventricular walls at this stage was shown in **Figure 1F**. Consistently, the atrial trabeculae were almost absent in the *Tie1* mutant mice at E14.5, while trabeculae became sparse in both left and right ventricles (**Figure 1G-I**). There was also a trend of decrease in the coronary vessel density upon TIE1 deficiency (**Figure 1J**). By the later stages of embryonic development (E17.5), TIE1 deficiency resulted in more severe cardiac defects, including a significant reduction in heart size, absence of atrial trabeculae, and disorganized trabecular structures in ventricles, together with a decrease in the ventricular wall thickness as well as the coronary vessel density (**Figure 2A-F**). The defective atrial trabecula formation was also validated in another *Tie1* mutant line targeting its intracellular kinase domain (*Tie1*^Δ*ICD/*Δ*ICD*^, **Figure 2–figure supplement 1**A-E). Consistent with the above results, TIE1 deficiency resulted in the lack of atrial trabeculation while trabeculae in ventricles became sparse in *Tie1* mutants at E14.5.

**Figure 2.**
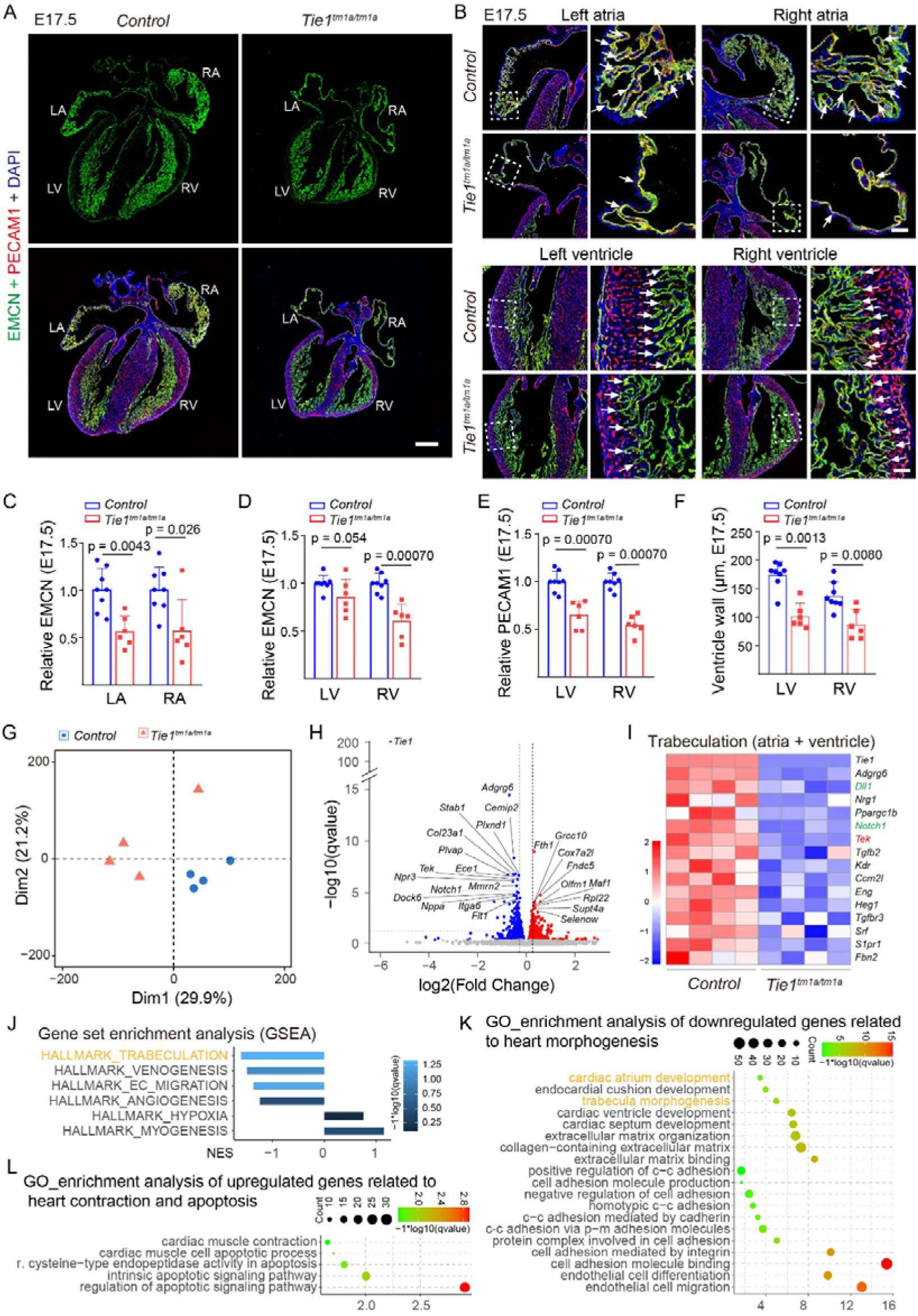
Suppression of cardiac trabeculation-related gene expression by TIE1 deficiency. **A-B**. Analysis of heart trabeculae of *Tie1^tm1a/tm1a^* and littermate controls by immunostaining for EMCN (green), PECAM1 (red) and DAPI (blue) at E17.5. Note that the ventricular trabeculae of *Tie1^tm1a/tm1a^* mice become sparse compared with those of control mice (white arrows). Arrows point to the trabecula-associated endocardium. **C-F**. Quantification of the EMCN-positive area in the atrial and ventricular trabeculae (**C**, LA: *Tie1^tm1a/tm1a^*: 0.56±0.16, n=6; *Control*: 1.00±0.22, n=8; *P*=0.0043. RA: *Tie1^tm1a/tm1a^*: 0.57±0.33, n=6; *Control*: 1.00±0.24, n=8; *P*=0.026. **D**, LV: *Tie1^tm1a/tm1a^*: 0.86±0.18, n=6; *Control*: 1.00±0.082, n=8; *P*=0.054. RV: *Tie1^tm1a/tm1a^*: 0.61±0.18, n=6; *Control*: 1.00±0.10, n=8; *P*=0.00070). Quantification of the PECAM1-positive area in ventricular walls (**E**, LV: *Tie1^tm1a/tm1a^*: 0.66±0.14, n=6; *Control*: 1.00±0.11, n=8; *P*=0.00070. RV: *Tie1^tm1a/tm1a^*: 0.55±0.10, n=6; *Control*: 1.00±0.088, n=8; *P*=0.00070.). Quantification of the ventricular wall thickness (**F**, LV: *Tie1^tm1a/tm1a^*: 101.97 ± 22.44 µm, n=6; *Control*: 174.09 ± 22.84 µm, n=8; *P*=0.0013. RV: *Tie1^tm1a/tm1a^*: 87.14±26.20 µm, n=6; *Control*: 137.35±24.06 µm, n=8; *P*=0.0080.) at E17.5. Except the ventricular wall thickness, the quantification data of mutant mice was normalized against that of littermate control mice. **G-L**. RNA-seq analysis of whole hearts of *Tie1^tm1a/tm1a^*and littermate controls (E14.5, n=4 per group). **G**. PCA analysis showed that *Tie1^tm1a/tm1a^* and *control* groups were divided into two clusters. **H**. Volcano plot visualization of the differentially expressed genes (DEGs, P-value ≤ 0.05, |Log2Foldchange| > 0). **I**. Heatmap of significantly downregulated cardiac trabeculation genes (P-value ≤ 0.05). **J**. GSEA analysis showed the decrease of enrichment in trabeculation, venous development, and endothelial cell migration pathways. **K-L**. GO analysis revealed that gene related to cardiac chamber morphogenesis and trabeculae development, extracellular matrix, cell adhesion and endothelial migration/differentiation were significantly downregulated (**K**), while cardiac contraction and apoptosis related genes were significantly upregulated (**L**). In C-F, statistical analysis was performed using the unpaired nonparametric Mann-Whitney U test. Values are represented as means ± SD of at least three independent technical replicates. LA: Left atria, RA: Right atria, LV: Left ventricle, RV: Right ventricle; Scale bar: 400 μm in A, 50 μm in B; c-c: cell-cell, p-m: plasma membrane in K.

There was also a trend of decrease in ventricular wall thickness as well as the coronary vessel density in the *Tie1*^Δ*ICD/*Δ*ICD*^ mice. Consistently, the atrial trabecular defects could also be reflected by the fractal analysis of cardiac trabecular complexity in atria from *Tie1* mutant and control mice at E14.5 and E17.5 using the Fraclac Plugin in ImageJ, while the relatively minor defects in the ventricles were not detected by the method (*Tie1^tm1a/tm1a^*, **Figure 2–figure supplement 2**; Supplement Table 2).

Consistent with the previous observation [25], the lack of TIE1 also disrupted the coronary vein formation at the later embryonic stages (E14.5 and E17.5), including the decrease of vein diameter and the increase of vein-associated angiogenic sprouting (**Figure 2–figure supplement 3**A-D).

### Alteration of atrial versus ventricular gene regulatory networks upon TIE1 deficiency

To explore the mechanism underlying TIE1 in heart development, we performed the RNA sequencing (RNA-seq) analysis of hearts to examine the alteration of cardiac transcriptome upon TIE1 deficiency. Lack of TIE1 led to a dramatic change in the overall transcriptional profile (**Figure 2G-L**), including the suppression of gene expression related to the endocardial / endothelial development and cardiac chamber morphogenesis, such as *Tek* (normalized by *Pecam1*, *Tie1^tm1a/tm1a^*: 0.89 ± 0.031, n=4; *WT*: 1.00 ± 0.12, n=4; *P* = 0.057.). Specifically, pathways related to extracellular matrix, cell adhesion and endothelial cell migration were downregulated (**Figure 2K**), while the cardiac contraction and cell apoptosis pathways were upregulated (**Figure 2L**). This was further confirmed by the GSEA analysis using the custom gene sets (**Figure 2J**). The gene sets included the following entries based on the GO database (HALLMARK_TRABECULATION, HALLMARK_EC_MIGRATION) and the fifty-three entries from the previous publication [25]. Consistent with previous reports [25], TIE1 deficiency led to the downregulation of venous genes. In addition, key regulatory factors for cardiac trabecular development were downregulated, including *Adgrg6* (adhesion G protein-coupled receptor G6), *Nrg1* (neuregulin 1), and *Notch1* (Notch receptor 1) and *Dll1* (Delta Like Canonical Notch Ligand 1). Notably, we found that the expression of the TIE receptor family member *Tek* was also significantly reduced (**Figure 2I**), suggesting that TIE1 acts, at least partially, via TIE2 in the regulation of cardiac trabecular morphogenesis.

To better visualize the trabecular structures, particularly the atrial internal muscular network, we performed the whole-mount immunostaining of the hearts at the four-chambered stage. As shown in **Figure 3A**, TIE1 deficiency led to the absence of the atrial trabecular network, with minor defects in the ventricles including a trend of sparse trabeculae (the 3D views shown in **Figure 3–figure supplement 1**). To further investigate the differential requirement of TIE1 in atrial and ventricular trabeculation, we collected atria and ventricle tissues separately from *Tie1* mutant and control mice (E14.5) for the RNA-seq analysis. We found that the number of differentially expressed genes was greater in atria than in the ventricles after the loss of TIE1 (**Figure 3B-G**). Similar to the results from the whole heart RNA-seq analysis, the transcript levels of genes related to atrial development were decreased. Although a similar trend of decrease in genes involved in the cardiac morphogenesis was also observed in the ventricles, the enrichment levels and the number of genes involved were much lower in ventricles than in atria. Specifically, 17 trabecula-related genes were downregulated in the atria, while there was only 8 in the ventricles. Notably, *Tek* and *Dll1* were downregulated in both the atria and ventricles while *Notch1* only in atria of *Tie1* mutant mice (**Figure 3F-G; Figure 2–figure supplement 1**F-G; Supplement Table 3 and Supplement Table 4). Interestingly, the expression of endothelial *Tie1*, *Tek* and *Notch1* was higher in atria than in ventricles (**Figure 3H, Figure 1–figure supplement** 1D, Supplement Table 5). The top GO terms enriched for the up- or down-regulated genes were related to the biological processes of heart development, including those in trabecular formation, extracellular matrix organization and binding, intercellular adhesion and endothelial cell migration in both atria and ventricles (**Figure 3I-J**). In addition, the analysis of scRNA-seq data for the ligand-receptor interactions showed that the atrial_EC (endocardial cells) and atrial_CM (cardiomyocytes) exhibited stronger ANGPT1-TIE2 signaling compared with those in ventricles (**Figure 1–figure supplement** 1E). This may account for the differential requirement of TIE1 and also TIE2 in the process of atrial versus ventricular trabeculation.

**Figure 3.**
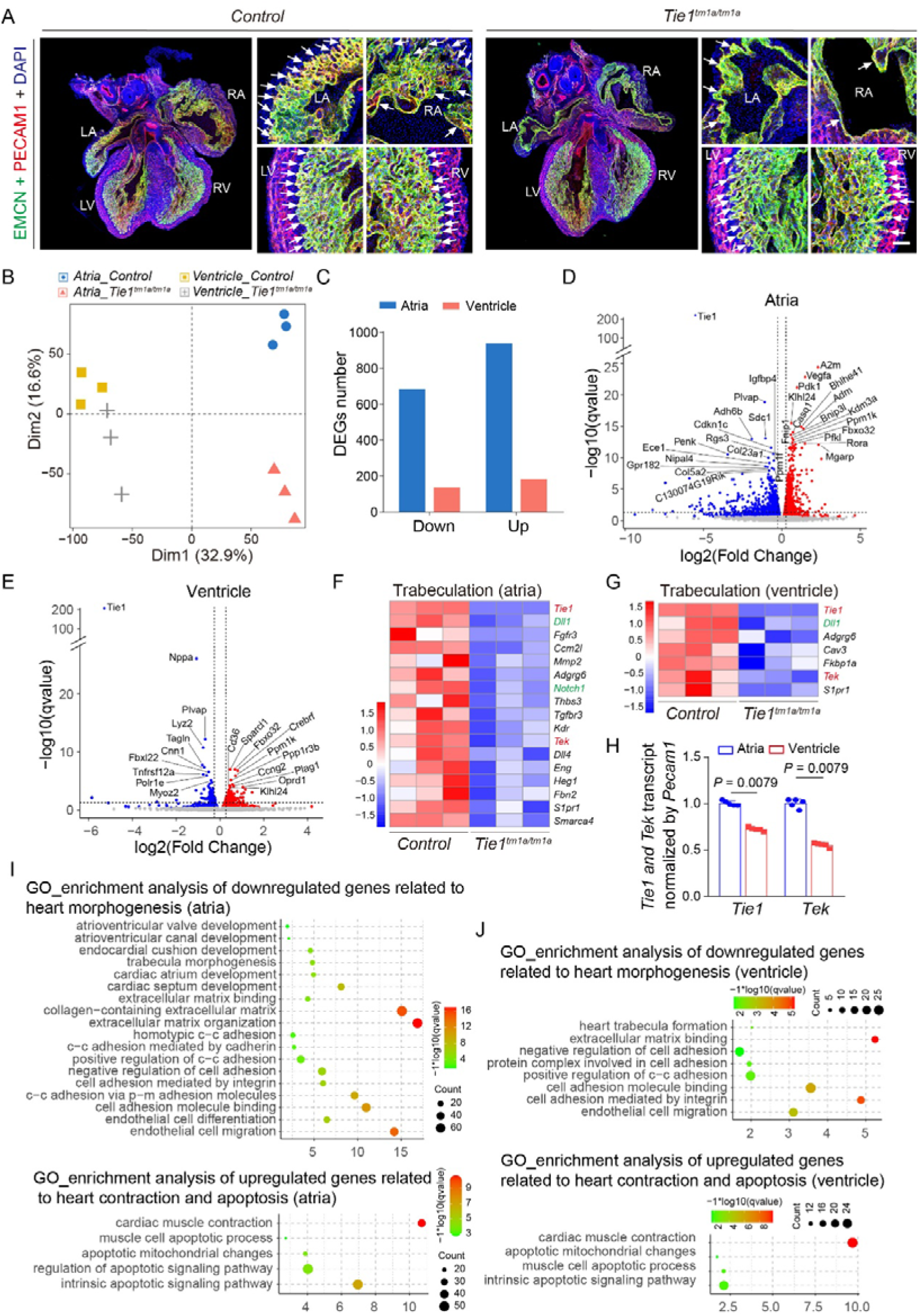
Differential effects of *Tie1* deficiency on atrial versus ventricular trabeculation related gene networks. **A**. Visualization of heart trabeculae structure of *Tie1^tm1a/tm1a^*and littermate controls by whole-mount immunostaining for EMCN (green), PECAM1 (red) and DAPI (blue) at E14.5. Note that *Tie1* deletion resulted in the loss of atrial trabecular network with minor defects with ventricular trabecular structures (white arrows). Arrows point to the trabecula-associated endocardium. **B-J**. RNA-seq analysis of atrium and ventricles of *Tie1^tm1a/tm1a^* and littermate controls (E14.5, n=3 per group). **B**. PCA analysis results showed clear separation between the atria and ventricles of *Tie1^tm1a/tm1a^* and control mice. **C**. Differential expression analysis (P-value ≤ 0.05, |log2(Foldchange)| ≥ 0.15) identified over 693 down-regulated and 948 up-regulated atrial genes, while 144 down-regulated and 190 up- regulated genes in ventricles. **D-E**. Volcano plots of atrial (**D**) and ventricular (**E**) differential expressed genes (DEGs). **F-G**. Heatmap showed significantly downregulated (P-value ≤ 0.05) heart trabeculation genes in atrium (**F**) and ventricles (**G**), including *Tek*, *Dll1* and *Notch1*. **H**. Quantification of *Tie1* and *Tek* transcript levels (normalized by *Pecam1*) in atrium and ventricles of *Tie1^tm1a/tm1a^*and littermate controls (*Tie1*: Ventricles: 0.72 ± 0.021, n = 5; Atria: 1.00 ± 0.026; *P* = 0.0079. *Tek*: Ventricle: 0.55 ± 0.024; Atria: 1.00 ± 0.048; *P* = 0.0079). **I-J**. GO enrichment analysis of the DEGs in atria and ventricles including the heart trabecula morphogenesis, extracellular matrix, cell adhesion and endothelial migration related process down-regulated while genes related to the cardiac contraction, apoptosis process up-regulated. There was a more obvious enrichment detected in the atria. In H, statistical analysis was performed using the unpaired nonparametric Mann-Whitney U test. Values are represented as means ± SD of at least three technical replicates. LA: left atria, RA: right atria, LV: left ventricle, RV: right ventricle; Scale bar: 50 μm in A; c-c: cell-cell, p-m: plasma membrane in I-J.

### Stronger ANGPT1-TIE2 pathway interaction between atrial endocardium and myocardium

Using the published scRNA-seq data [34], we also analyzed the ligand-receptor interactions among endocardial cells and cardiomyocytes in atria and ventricles respectively during the embryonic stage (E10.5-E18.5). We found that the ANGPT1-TIE2 signaling between atrial cardiomyocytes and endocardial cells was stronger than that in ventricles (**Figure 1–figure supplement** 1E). To characterize temporal transcriptional changes, we analyzed the developmental expression profiles of representative genes in cardiac and endocardial cells from E10.5 to E18.5 using published single-cell RNA-sequencing (scRNA-seq) data. As shown in **Figure 1– figure supplement** 1F, *Angpt1* was higher across total atrial cardiac cells than those of ventricles, while there was no obvious difference with *Angpt2* and *Svep1* (left panel). Concurrently, the expression of *Tie1*, *Tek*, and *Npr3* was detected in atrial and ventricular endocardial cell populations (right panel), revealing a trend of higher Tie1 in atrial cells (right panel).Consistently, the crucial role of endothelial TIE2 in the atrial trabeculation was validated in this study by employing a conditional knockout mouse model targeting *Tek* (also known as *Tie2* gene; *Tek^Flox/-^; Cdh5-Cre^ERT2^*; named *Tek^iECKO^*). As the complete knockout of *Tek* gene leads to embryonic lethality before E10.5, we first performed the induced endothelial *Tek* deletion starting from E12.5 (**Figure 4A-B**). We found that TIE2 insufficiency led to an obvious decrease of trabeculae in the atrium 48 h later (E14.5), while there were no obvious defects observed in the ventricular trabeculae at this stage (**Figure 4C-E**). The coronary vessel density in the ventricular wall as well as the wall thickness, including left and right ventricles, were not obviously altered when examined 48 h later (**Figure 4F**). In separate experiments with the induced gene deletion from E12.5 to E14.5, embryos were collected for the analysis at 72 hours (E15.5) or later (E17.5). The formation of atrial trabeculae was completely missing as shown in **Figure 4A, G-H**. The defective trabecula formation was also detected in the ventricles, especially at later embryonic stages (E17.5, **Figure 4A, G-H**). TIE2 insufficiency also led to the defective formation of the coronary veins as previously shown in skin and mesentery (**Figure 2–figure supplement 3**E, G).

**Figure 4.**
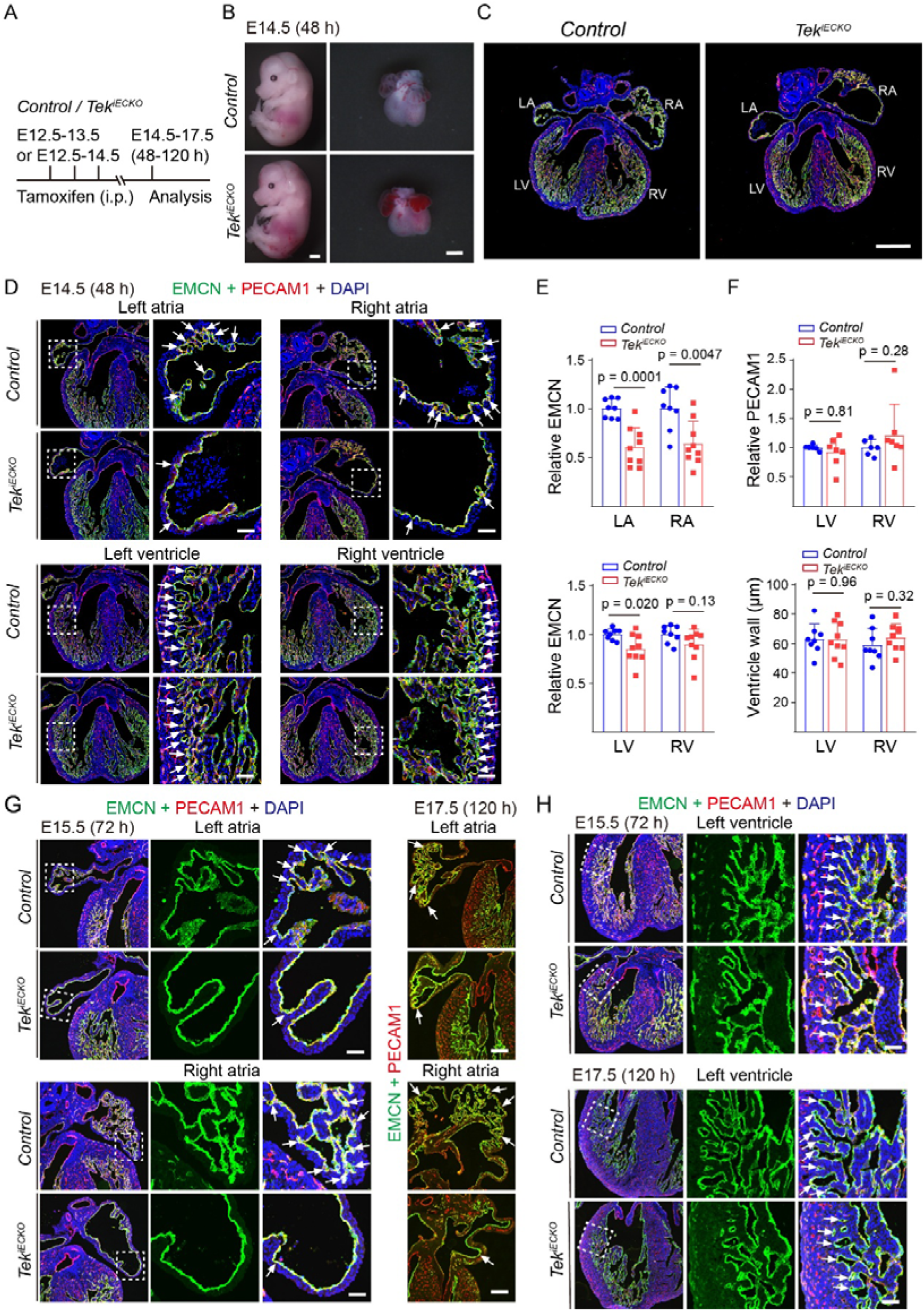
Impaired atrial trabecular formation upon TIE2 insufficiency. **A.** Tamoxifen intraperitoneal (i.p.) administration and analysis scheme. **B-D**. Analysis of heart trabeculae of *Tek^iECKO^*and littermate controls by immunostaining for EMCN (green), PECAM1 (red) and DAPI (blue) 48 hours later (tamoxifen treatment starting at E12.5). **E-F**. Quantification of the EMCN-positive area in the atrial and ventricular trabeculae (**E**, LA: *Tek^iECKO^*: 0.60±0.20, n=9; *Control*: 1.00±0.097, n=8; *P*=0.00010. RA: *Tek^iECKO^*: 0.64±0.23, n=9; *Control*: 1.00±0.22, n=8; *P*=0.0047. LV: *Tek^iECKO^*: 0.85±0.15, n=9; *Control*: 1.00±0.057, n=8; *P*=0.020. RV: *Tek^iECKO^*: 0.90±0.16, n=9; *Control*: 1.00±0.089, n=8; *P*=0.13.), quantification of the PECAM1-positive area in ventricular walls (**F**, LV: *Tek^iECKO^*: 0.92±0.25, n=7; *Control*: 1.00±0.048, n=6; *P*=0.81. RV: *Tek^iECKO^*: 1.22±0.53, n=7; *Control*: 1.00±0.13, n=6; *P*=0.28.) and quantification of the ventricular wall thickness (**F**, LV: *Tek^iECKO^*: 62.35±11.70 µm, n=9; *Control*: 62.59±10.26 µm, n=8; *P*=0.96. RV: *Tek^iECKO^*: 63.87± 9.19 µm, n=9; *Control*: 58.87±11.02 µm, n=8; *P*=0.32.) at E14.5. Except the ventricular wall thickness, the quantification data of mutant mice was normalized against that of littermate control mice. **G-H**. Analysis of heart trabeculae of *Tek^iECKO^* and littermate controls by immunostaining for EMCN (green), PECAM1 (red) and DAPI (blue) 72 hours after the induced gene deletion or later (tamoxifen treatment starting at E12.5). Arrows point to the trabecula-associated endocardium. In E-F, statistical analysis was performed using the unpaired t test. Values are represented as means ± SD of at least three independent technical replicates. LA: left atria, RA: right atria, LV: left ventricle, RV: right ventricle; Scale bar: 500 μm (embryo) and 400 μm (heart) in B, 400 μm in C, 50 μm in D, G, H.

### Functional synergy between TIE1 and TIE2 during atrial trabeculation

To investigate the synergistic effects of TIE1 and TIE2 in the regulation of cardiac morphogenesis, we generated a double knockout model targeting *Tie1* and *Tek* (*Tie1*^Δ*ICD/*Δ*ICD*^*; Tek^+/-^*). There were no obvious defects observed with the atrial or ventricular structures upon *Tie1* deficiency alone at E10.5 (**Figure 5A-B**). Intriguingly, loss of *Tie1* combined with the heterozygous deletion of *Tek* suppressed both atria and ventricle development (E10.5), including reduced heart size, abnormal atrial chamber formation, and fewer trabecular structures in the ventricles (**Figure 5C-D**), leading to embryonic lethality before E12.5.

**Figure 5.**
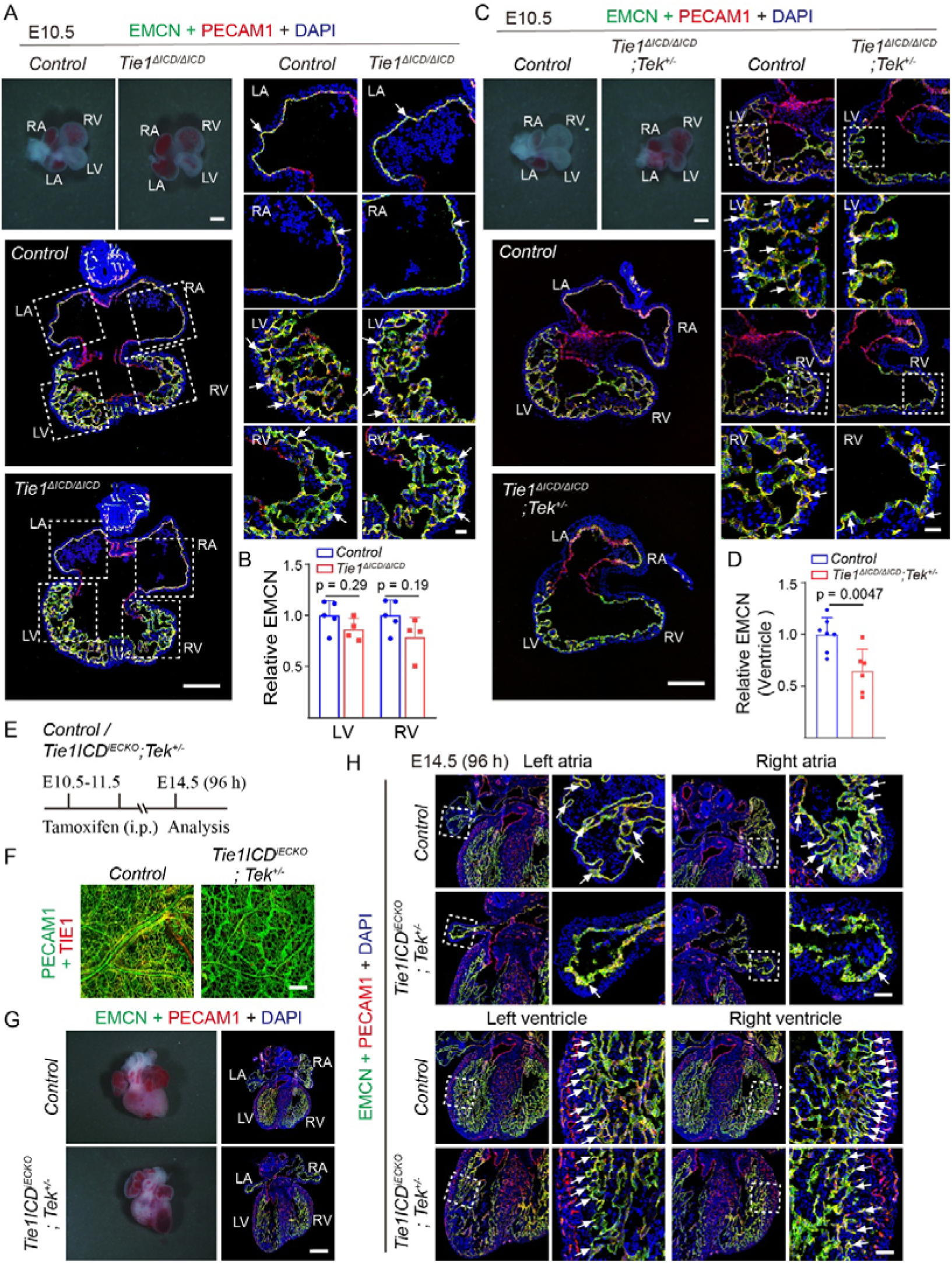
Requirement of TIE1 and TIE2 in atrial and ventricular chamber morphogenesis. **A-D.** Analysis of heart trabeculae of *Tie1*^Δ*ICD/*Δ*ICD*^, *Tie1*^Δ*ICD/*Δ*ICD*^*;Tek^+/-^*and littermate controls by immunostaining for EMCN (green), PECAM1 (red) and DAPI (blue) at E10.5. Note that there were no obvious defects observed with the atrial or ventricular structures upon *Tie1* deficiency alone at E10.5, while loss of *Tie1* combined with the heterozygous deletion of *Tek* suppressed both atria and ventricle development. White arrows point to the trabecular structures. Quantification of the EMCN-positive area in the ventricular trabeculae at E10.5, and the quantification data of mutant mice was normalized against that of littermate control mice (**B**, LV: *Tie1*^Δ*ICD/*Δ*ICD*^: 0.86±0.11, n=4; *Control*: 1.00±0.14, n=5; *P*=0.29. RV: *Tie1*^Δ*ICD/*Δ*ICD*^: 0.79±0.19, n=4; *Control*: 1.00±0.15, n=5; *P*=0.19. **D**, *Tie1*^Δ*ICD/*Δ*ICD*^*;Tek^+/-^*: 0.64±0.22, n=6; *Control*: 1.00±0.16, n=7; *P*=0.0047). **E**. For the induced gene deletion, tamoxifen intraperitoneal (i.p.) administration (starting from E10.5) and analysis scheme (at E14.5-E16.5). **F.** The deletion efficiency of TIE1 was examined by the immunostaining of skins from *Tie1ICD^iECKO^;Tek^+/-^*and littermate control mice at E16.5 for PECAM1 (green) and TIE1 (red). **G-H**. Analysis of heart trabeculae of *Tie1ICD^iECKO^;Tek^+/-^*mice and littermate controls by immunostaining for EMCN (green), PECAM1 (red) and DAPI (blue) at E14.5. Note that the atrial trabeculae structure was missing, while the ventricular trabeculae became sparse (white arrows). Arrows point to the trabecula-associated endocardium. In B and D, statistical analysis was performed using the unpaired nonparametric Mann-Whitney U test. Values are represented as means ± SD of at least three independent technical replicates. LA: left atria, RA: right atria, LV: left ventricle, RV: right ventricle. Scale bar: 200 μm in A(left) and C(left), 30 μm in A(right) and C (right), 400 μm in G, 200 μm in F and 50 μm in H.

To further explore the synergy of TIE1/TIE2 in the heart development at later stages of embryonic development, we generated an inducible mouse model with the endothelial deletion of *Tie1* combined with *Tek* heterozygous knockout (*Tie1ICD^Flox/-^ ;Cdh5-Cre^ERT2^;Tek^+/-^*, named *Tie1ICD^iECKO^;Tek^+/-^*). The induced gene deletion was performed by the tamoxifen treatment starting from E10.5, and mice were sacrificed for analysis after 96 hours (E14.5). The gene deletion efficiency was confirmed by the immunostaining for PECAM1 plus TIE1. TIE1 insufficiency combined with the heterozygous loss of TIE2 led to the suppression of atrial trabeculation, while the trabeculae at both right and left ventricles became sparse and irregular (**Figure 5E-H**). Furthermore, when the induced gene deletion was performed at E12.5, the defective atrial trabeculation were already detectable in the mutant mice 48 hours later (E14.5 *Tie1ICD^iECKO^;Tek^+/-^*; **Figure 6 A-D**). Consistent with the observation with the endothelial *Tek* deletion for 48 hours, there was no obvious defects with the ventricular trabeculation, coronary vessel density and ventricular wall thickness (*Tie1ICD^iECKO^;Tek^+/-^*; **Figure 6D-E**, Table 2). The disruption of cardiac trabeculation became more obvious when analyzed 72-96 hours later, particularly in atria but less severe with ventricles (*Tie1ICD^iECKO^; Tek^+/-^,* E15.5-E16.5; **Figure 6 F-G**, Table 3). There was also a trend of decrease in coronary vessel density and ventricular wall thickness as shown at later stages (E16.5, **Figure 6 F, H**, Table 3). In addition, the TIE1/TIE2 insufficiency also resulted in the disruption of cardiac vein development as shown in (**Figure 2–figure supplement 3**F, H), as previously reported in skin and retina [25].

**Figure 6.**
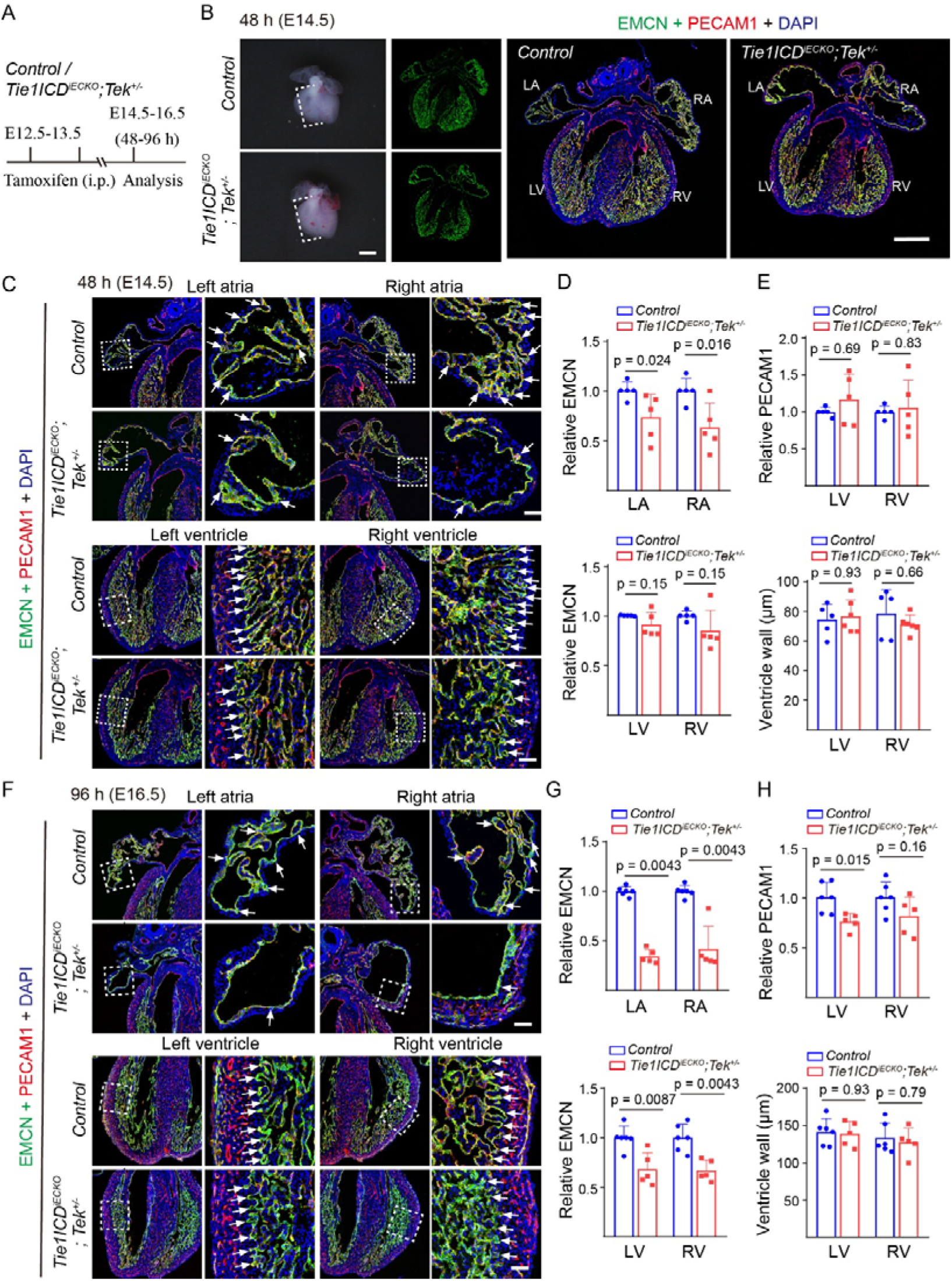
Synergistic roles of TIE1 and TIE2 in atrial trabeculation. **A**. Tamoxifen intraperitoneal (i.p.) administration and analysis scheme. **B-H**. The induced gene deletion by tamoxifen started at E12.5. Analysis of heart trabeculae of *Tie1ICD^iECKO^;Tek^+/-^*and littermate controls by immunostaining for EMCN (green), PECAM1 (red) and DAPI (blue) at E14.5 (**B-C**) and E16.5 (**F**). Note that the atrial trabeculae were absent and the ventricular trabeculae were slightly sparse in the double mutant mice at E14.5 (**B-C**), and the phenotype became more severe at E16.5 (**F**). Quantification of the EMCN-positive area in the atrial and ventricular trabeculae (**D**, E14.5; **G**, E16.5), the PECAM1-positive area in ventricular walls and the ventricular wall thickness (**E**, E14.5; **H**, E16.5). Arrows point to the trabecula-associated endocardium. The quantification data of mutant mice was normalized against that of littermate control mice as shown in Table 2 and Table 3. Arrows in C and F point to the trabecula-associated endocardium. In D, E, G and H, statistical analysis was performed using the unpaired nonparametric Mann-Whitney U test. Values are represented as means ± SD of at least three independent technical replicates. LA: left atria, RA: right atria, LV: left ventricle, RV: right ventricle. Scale bar: 400 μm in B, 50 μm in C and F.

**Table 2.** Quantification of the trabecular area (EMCN^+^), blood vessel density in ventricular wall (PECAM1^+^) and ventricular wall thickness in *Tie1ICD^iECKO^;Tek^+/-^* and littermate controls (at E14.5, 48 hours after the induced gene deletion). Statistical analysis was performed using the unpaired nonparametric Mann-Whitney U test. Values are represented as means ± SD of at least three independent technical replicates.

|  |  | LA | RA | LV | RV |
| --- | --- | --- | --- | --- | --- |
| Relative EMCN | Control<br>(n=5, mean±sd) | 1.00±0.086 | 1.00±0.12 | 1.00±0.0050 | 1.00±0.048 |
|  | *Tie1*CD<sup>IECKO</sup>;Tek<sup>+/-</sup><br>(n=5, mean±sd) | 0.73±0.23 | 0.63±0.24 | 0.91±0.12 | 0.85±0.21 |
|  | *P* | 0.024 | 0.016 | 0.15 | 0.15 |
| **Relative**<br>**PECAM1** | *Control*<br>(n=5, mean±sd) | - | - | 1.00±0.060 | 1.00±0.083 |
|  | *Tie1*CD<sup>IECKO</sup>;Tek<sup>+/-</sup><br>(n=5, mean±sd) | - | - | 1.17±0.34 | 1.06±0.38 |
|  | *P* | - | - | 0.69 | 0.83 |
| **Ventricle**<br>**wall (μm)** | *Control*<br>(n=5, mean±sd) | - | - | 74.48±10.62 | 78.56±16.48 |
|  | *Tie1*CD<sup>IECKO</sup>;Tek<sup>+/-</sup><br>(n=6, mean±sd) | - | - | 76.87±11.33 | 71.33±6.25 |
|  | *P* | - | - | 0.93 | 0.66 |

**Table 3.** Quantification of the trabecular area (EMCN^+^), coronary vessel density in ventricular wall (PECAM1^+^) and ventricular wall thickness in *Tie1ICD^iECKO^;Tek^+/-^*and littermate controls 96 hours after the induced gene deletion (E16.5; tamoxifen treatment from E12.5-E13.5). Statistical analysis was performed using the unpaired nonparametric Mann-Whitney U test. Values are represented as means ± SD of at least three independent technical replicates.

|  |  | **LA** | **RA** | **LV** | **RV** |
| --- | --- | --- | --- | --- | --- |
| **Relative**<br>**EMCN** | *Control*<br>(n=6, mean±sd) | 1.00±0.048 | 1.00±0.059 | 1.00±0.12 | 1.00±0.14 |
|  | *Tie1*CD<sup>IECKO</sup>;Tek<sup>+/-</sup><br>(n=5, mean±sd) | 0.34±0.069 | 0.42±0.23 | 0.68±0.17 | 0.67±0.11 |
|  | *P* | 0.0043 | 0.0043 | 0.0087 | 0.0043 |
| **Relative**<br>**PECAM1** | *Control*<br>(n=6, mean±sd) | - | - | 1.00±0.15 | 1.00±0.16 |
|  | *Tie1*CD<sup>IECKO</sup>;Tek<sup>+/-</sup><br>(n=5, mean±sd) | - | - | 0.76±0.083 | 0.81±0.19 |
|  | *P* | - | - | 0.015 | 0.16 |
| **Ventricle**<br>**wall (μm)** | *Control*<br>(n=6, mean±sd) | - | - | 141.45±18.02 | 133.36±20.44 |
|  | *Tie1*CD<sup>IECKO</sup>;Tek<sup>+/-</sup><br>(n=5, mean±sd) | - | - | 138.90±16.66 | 127.81±18.81 |
|  | *P* | - | - | 0.93 | 0.79 |

### Requirement of TIE receptors during postnatal atrial trabecular remodeling

The role of TIE1 and its synergy with TIE2 in cardiac trabecular development was further confirmed at the neonatal stage (*Tie1ICD^iECKO^* or *Tie1ICD^iECKO^; Tek^+/-^*) with the induced gene deletion from postnatal day 1 (P1-4) and the analysis performed at P21. The *Tie1* deletion was confirmed by the vascular defects in retinas of *Tie1ICD^iECKO^* or *Tie1ICD^iECKO^; Tek^+/-^* mice as previously reported (**Figure 7B, E**) [25]. TIE1 insufficiency alone had no obvious impact on the heart weight and also the atrial or ventricular trabeculation analyzed within three weeks (**Figure 7 A-D**). Consistent with the studies at embryonic stages, the induced postnatal gene deletion (*Tie1ICD^iECKO^;Tek^+/-^*) showed the retarded heart development by the quantification of heart-body weight (**Figure 7 F, G**), and showed abnormal atrial trabeculation while the trabeculae in the ventricles had no obvious defects (**Figure 7F, H**). There was a significant decrease of trabecular structures (white arrows in **Figure 7H**) in atria of *Tie1ICD^iECKO^; Tek^+/-^* mice compared with that of the control mice (LA, *Tie1ICD^iECKO^; Tek^+/-^*: 0.69±0.16, n=11; *Control*: 1.00±0.28, n=16; *P*=0.0022. RA, *Tie1ICD^iECKO^; Tek^+/-^*: 0.77±0.15, n=11; *Control*: 1.00±0.11, n=16; *P*=0.0002). The effects of TIE1 and TIE2 deficiency or insufficiency on heart trabecular morphogenesis were summarized in **Figure 7I**. Based on the previous findings about the TIE1/TIE2 in venous EC specification [25, 28], we propose that TIE1 acts in synergy with TIE2 in the specification of atrial endocardial cells (**Figure 7I**). TIE1 deficiency or TIE1/TIE2 insufficiency results in a more severe defects with the atrial trabeculation, suggesting the differential requirement of TIE1 and TIE2 in the endocardium-mediated organization of the atrial internal muscular network.

**Figure 7.**
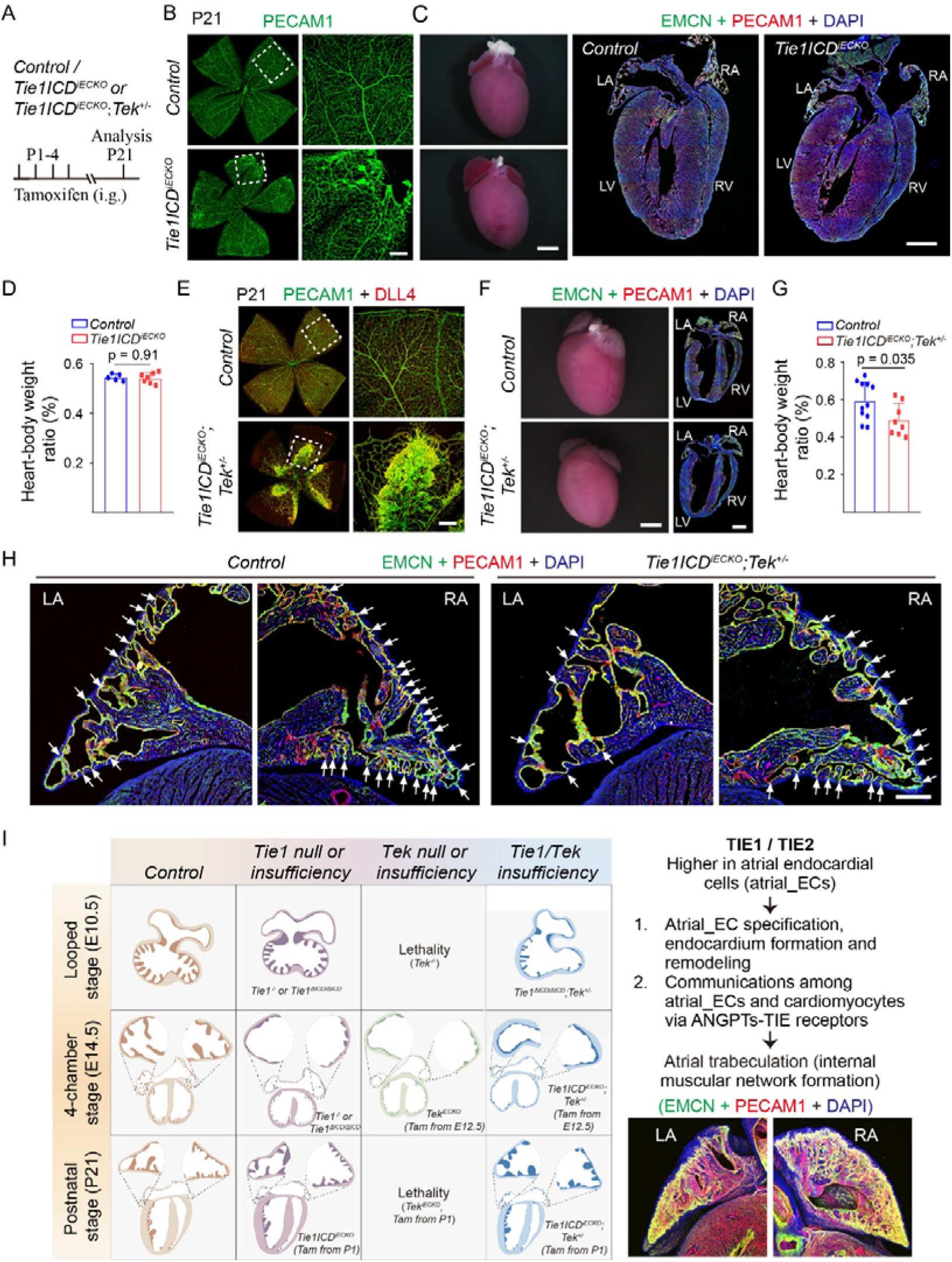
Abnormal atrial trabecular remodeling after the postnatal deletion of *Tie1* combined with one null allele of *Tek*. **A.** Tamoxifen intragastric (i.g.) administration and the analysis scheme. **B-H.** Analysis of blood vessels in the retinas of *Tie1ICD^iECKO^* (**B**), *Tie1ICD^iECKO^;Tek^+/−^*(**E**) and *control* mice by whole-mount immunostaining for PECAM1 (green), DLL4 (red) at P21. Note that hemangioma-like vascular tufts observed in *Tie1ICD^iECKO^* and *Tie1ICD^iECKO^;Tek^+/−^*mice. The retinal vascular defects were used to confirm the efficiency of induced gene deletion. Analysis of heart trabeculae in *Tie1ICD^iECKO^* (**C**), *Tie1ICD^iECKO^;Tek^+/−^*(**F**) and control mice by the immunostaining for EMCN (green), PECAM1 (red) and DAPI (blue) at P21. TIE1 insufficiency alone had no obvious effect on heart size (**D,** *Tie1ICD^iECKO^*: 0.54±0.021, n=7; *Control*: 0.54±0.014, n=5; *P*=0.91.) and atrial trabecular structures (**C**). The *Tie1ICD^iECKO^;Tek^+/-^*mice displayed smaller hearts (**G**, *Tie1ICD^iECKO^;Tek^+/-^*: 0.49±0.086, n=8; *Control*: 0.59±0.10, n=11; *P*=0.035.) and abnormal atrial trabecular morphology (**F,H**). White arrows in H point to the trabecula-associated endocardium in the atria. **I.** Schematic illustration of TIE1, TIE2 and their synergy in cardiac chamber morphogenesis. TIE1 deficiency (*Tie1^tm1a/tm1a^*, *Tie1*^Δ*ICD/*Δ*ICD*^), TIE2 (*Tek^iECKO^*) or TIE1/TIE2 insufficiency (*Tie1*^Δ*ICD/*Δ*ICD*^*;Tek^+/-^* or *Tie1ICD^iECKO^;Tek^+/−^*) results in a more severe defect with the atrial trabeculation than that of ventricles. Based on the previous findings about the TIE1/TIE2 in venous EC specification [25, 28] and findings from this study, we propose that TIE1 acts in synergy with TIE2 in the specification and remodeling of atrial endocardium, and that there is a differential requirement of TIE receptors in atrial versus ventricular wall morphogenesis. In D and G, statistical analysis was performed using the unpaired nonparametric Mann- Whitney U test (D) and the unpaired t test (G). Values are represented as means ± SD of at least three independent technical replicates. Scale bar: 50 μm in B and E, 1 mm in C and F, 200 μm in H.

## Discussion

Developmental defects or pathological remodeling of atria may result in the atrial dysfunction leading to the heart failure [39]. In contrast to the better understanding of ventricular chamber morphogenesis, the dynamic process and regulatory networks of the atrial internal muscular network formation (known as atrial trabeculation) are inadequately elucidated. TIE1 variants or loss of function mutations were reported in a subset of lymphedema patients [32, 33], and it is unknown whether the patients have also cardiac defects in addition to the lymphatic abnormality. We show in this study that the atrial trabeculation occurred later than in ventricles. Based on the analysis of the published scRNA-seq data by Feng et al. [34] and the bulk RNA-seq analysis of atria and ventricles separately in this study, we showed that *Tie1* and its homologue *Tek* were highly expressed in atrial endocardial cells than those of ventricles. By employing the genetically modified mouse models targeting *Tie1*, *Tek* and both, we found that TIE1 was differentially required in the process of atrial versus ventricular trabeculation and acted in synergy with TIE2 in the process of cardiac trabeculation, especially in atria.

### Endocardial versus vascular effects of TIE1

Cardiac chamber morphogenesis involves the coordinated endocardial-myocardial interaction governed by a plethora of cardiovascular regulators [1]. It is worth noting that most of the reported endothelial regulators have important roles in both heart and vascular development. ANGPT1 has been shown to participate in the regulation of atria and cardiac vein development [18, 40]. We have demonstrated that TIE1 and TIE2 are required for the vein specification via regulating the protein stability of COUP-TFII [28]. Disruption of COUP-TFII led to the defective atrial and venous formation [29, 31]. Defective cardiac venous formation was also observed in this study in *Tie1* or *Tek* mutant mice at later embryonic stages. It is therefore important to dissect whether cardiac defects in the mutants are a direct effect or secondary to the coronary vascular defects. By performing a serial analysis of the developing hearts during embryogenesis (including E10.5, E12.5-, E14.5, E16.5, E17.5), we found in this study that TIE1 deficiency resulted in an impaired atrial trabeculation readily detectable at embryonic day 12.5 (E12.5), while there were only minor defects observed with the ventricular trabeculae at the same stage. In addition, *Tek* deficiency leads to the embryonic lethality before E10.5 [28, 41]. Interestingly, the defective atrial trabeculae formation was detected within 48 hours after the induced deletion of endothelial *Tek* (starting from E12.5), while there were no obvious defects with coronary vessels at the same stage. This was further confirmed in the *Tie1/Tek* double mutant mice with defective atrial trabeculation detected within 48 hours after the induced deletion of *Tie1* combined with one null allele of *Tek* (*Tie1ICD^iECKO^;Tek^+/-^*). Lack of TIE1 did not produce obvious developmental defects in mutant embryos by E10.5 as previously reported [25, 27, 28]. Therefore, the differential requirement of TIE1 may be related to the specification of atrial versus ventricular endocardium, which involves a synergistic role of TIE2, as previously reported in vein development [25]. Based on the above findings, it is likely that TIE1 and TIE2 are directly required for the endocardial cell-mediated organization of atrial trabecular structures. In addition, defects with the ventricular trabeculation and a trend of thinner ventricular walls were also observed at later stages after the *Tie1* deficiency or induced *Tie1/Tek* insufficiency. The cardiac trabeculation contributes to the thickening and maturation of the ventricular wall by being incorporated into the compact myocardium [4, 8]. It has been reported that endocardial cells contribute to the formation of coronary vascular network formation [2]. The defective ventricular wall morphogenesis may also be related to the decreased coronary vessel growth in Tie1 or *Tie1/Tek* mutant mice as observed in this study.

### Synergy of TIE1 and TIE2 in cardiac trabeculation

Loss of TIE1 alone did not produce an obvious defect with atrial chamber formation at initial stages of heart development (e.g. E10.5). However, the loss of *Tie1* combined with the heterozygous deletion of *Tek* disrupted the atrial and ventricular development at the same stage (E10.5). The synergy of TIE1 and TIE2 in heart development was further verified at later embryogenesis as well as the postnatal studies. The defects with the atrial trabeculation were already detectable 48 hours later in the induced knockout of *Tie1* plus the heterozygous *Tek* deletion (starting at E12.5, *Tie1ICD^iECKO^; Tek^+/-^*). This was further confirmed at the neonatal with the induced gene deletion from postnatal day 1 and the analysis 3-weeks later stage (*Tie1ICD^iECKO^; Tek^+/-^*), while TIE1 insufficiency alone (*Tie1ICD^iECKO^*) had no obvious impact on the atrial or ventricular trabecula development. TIE1 and TIE2 could form heterodimers, participate in the formation of endothelial cell-cell adherens junction as well as the EC migration and survival [19, 20, 42, 43]. Furthermore, TIE1 and TIE2 act in a synergistic manner to specify the vein formation via regulating the protein stability of the venous factor COUP-TFII [25]. As COUP-TFII is crucial for the atrial development [29, 31], this could provide some insights into the synergy of TIE1/TIE2 in the regulation of cardiac trabeculation especially in atria, likely involving COUP-TFII-mediated regulation of the atrial endocardial cell specification.

### Differential requirement of TIE receptors in atria and ventricles

So far, little is known about mechanisms underlying the differential regulation of atrial versus ventricular endocardial development. We found in this study that *Tie1* and *Tek* were expressed at higher levels in atrial endocardial cells than those of ventricles by the RNA-seq analysis at the single cell level or the separate analysis of atria and ventricles. By the analysis of scRNA-seq data from embryonic hearts for the ligand-receptor interactions, we showed that the atrial endocardial cells and cardiomyocytes exhibited stronger ANGPT1-TIE2 signaling compared with those in ventricles. This suggests that the differential requirement of TIE1 and TIE2 in cardiac chamber morphogenesis may, at least partly, be related to the differential expression levels of TIE receptors in atrial versus ventricular endocardial cells. In addition, we also found that *Dll1* was mainly expressed by endocardial cells from the analysis of single cell RNA-seq data [34], and that *Dll1* was among the most downregulated genes in the hearts from *Tie1* deficient mice. A similar trend of decreased expression with *Notch1* was also detected in atria but not in ventricles of *Tie1* deficient mice. As NOTCH1 signaling promotes the extracellular matrix degradation during the formation of endocardial projections that are critical for individualization of trabecular units [12, 14], this may also contribute to the differential requirement of TIE1 as well as TIE2 in atrial versus ventricular trabeculation. However, it is worthing pointing out that using a zebrofish model, Albu et al. reported no evidence for Nrg/ErbB or Notch signaling in atrial muscle network formation [44]. In addition to the decrease in *Notch1* as well as *Erg1* transcripts in *Tie1* mutants, we detected an increase in *Vegfa* transcript levels, which may result from compensatory changes in cardiac angiogenesis induced by hypoxic stress upon TIE1 deficiency. The biological significance of these transcriptomic changes requires further investigation.

In summary, the findings from this study demonstrate that endothelial TIE1 is crucial for atrial trabecular formation and plays a synergistic role with TIE2 during cardiac chamber morphogenesis. *Tie1* deficiency resulted in a significant downregulation of *Tek*, *Dll1* and *Notch1*, suggesting that the differential requirement of TIE receptors in atrial versus ventricular trabeculation may involve the DLL1-NOTCH1 pathway. While the precise mechanisms underlying TIE receptors in this chamber-specific regulation remain to be investigated, the dataset provides a valuable foundation for future mechanistic studies. Fully elucidating its downstream mechanism may require the isolation of primary endocardial cells for further study. However, our previous work using HUVECs have demonstrated that TIE1 acts synergistically with its homologue TIE2 to regulate AKT-mediated COUP-TFII protein stability [25, 28]. Additionally, utilizing single-cell RNA-seq to holistically map endocardial /endothelial specification in control versus *Tie1/Tek* mutant embryos also represents an exciting avenue for providing deeper mechanistic insights. Ultimately, expanding our knowledge in these directions will facilitate the development of novel interventions for atria-related cardiomyopathies.

## Supporting information

Figure 3-figure supplement 1 Control

Figure 3-figure supplement 1 Tie1tm1a

## Acknowledgements

The authors thank the staff in Animal facility of Soochow University for technical assistance.

## Data availability

The raw and processed RNA-seq data generated in this study have been deposited in the NCBI Gene Expression Omnibus (GEO) database (Accession numbers: GSE343212, GSE343213) and made publicly available upon publication.

---

**Figure 1–figure supplement 1.**
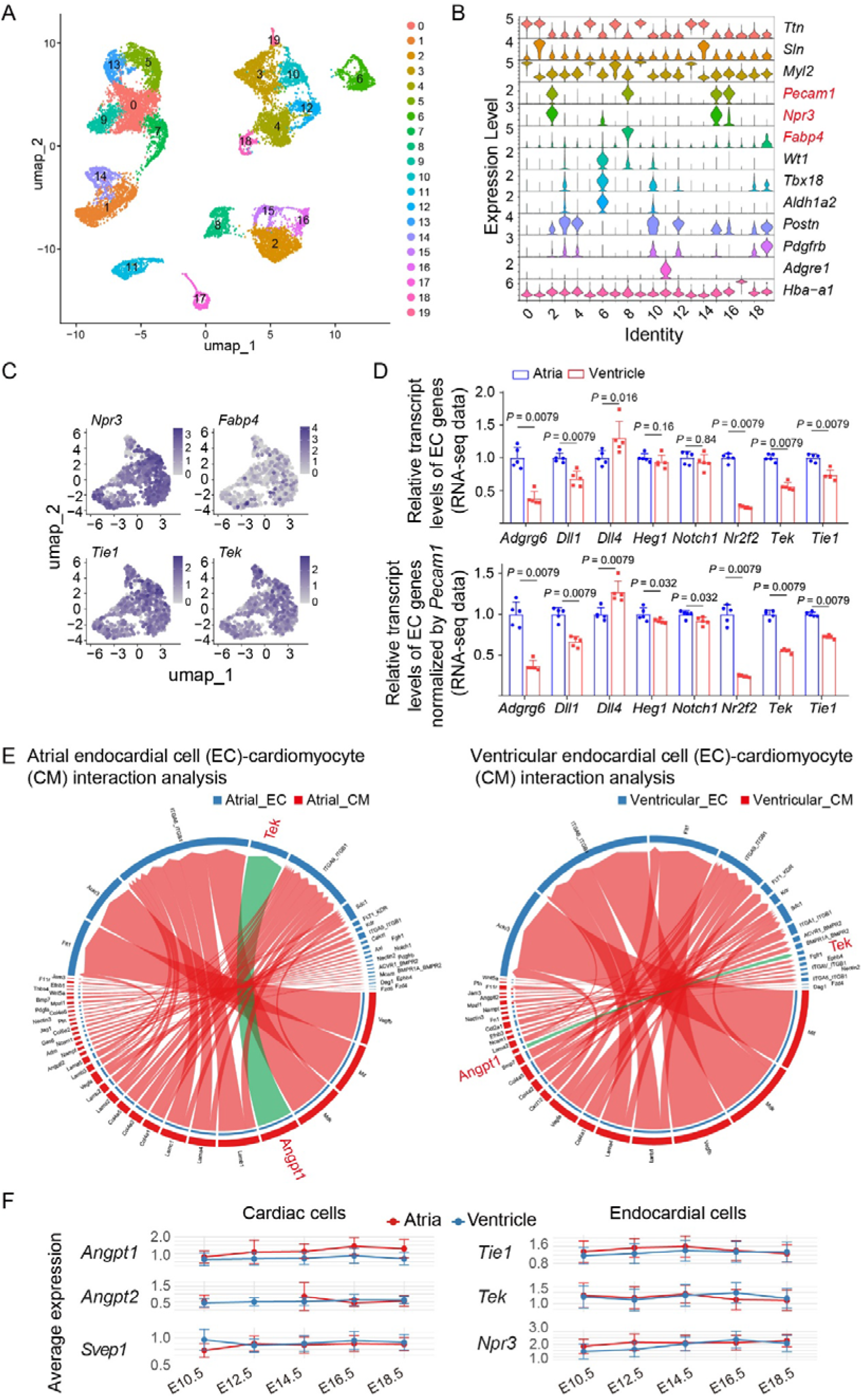
The scRNA-seq analysis of developmental changes of gene expression in the murine cardiac cells. **A-C.** Analysis of the single-cell RNA-seq data of embryonic hearts at E10.5 to E18.5 by Feng et al. Nat Commun 2022 (DOI: 10.1038/s41467-022-35691-7). UMAP visualization of 20 distinct cell clusters in the murine hearts during embryogenesis (E10.5 to E18.5, A). Violin plot shows the marker genes of all the clusters. Clusters 2, 15, and 16 were identified as endocardial cells (*Npr3*⁺). Marker genes for all clusters (B) are detailed in the Methods. Both *Tie1* and *Tek* are highly expressed in the endocardial cell population (*Npr3*⁺) while the coronary EC gene *Fabp4* is lowly expressed as shown in feature plots (C). **D**. Differential expression of trabeculation-related genes in atria versus ventricles at embryonic stage E14.5 (based on the RNA-seq analysis of atria and ventricles separately). The quantitative results are shown in Supplemental Table 5. **E**. Analysis of ligand-receptor interactions among endocardial cells and cardiomyocytes in atria and ventricles respectively during the embryonic stage (E10.5-E18.5). Note that the ANGPT1-TIE2 signaling between atrial cardiomyocytes and endocardial cells was stronger than that in ventricles. In D, statistical analysis was performed using the unpaired nonparametric Mann-Whitney U test. Values are represented as means ± SD of at least three technical replicates. **F**. Developmental expression profiles of representative genes in cardiac and endocardial cells (E10.5 to E18.5). Expression analysis was performed using published single-cell RNA-sequencing (scRNA-seq) data. (Left) Average expression levels of *Angpt1*, *Angpt2*, and *Svep1* across total atrial and ventricular cardiac cells. (Right) Average expression levels of *Tie1*, *Tek*, and *Npr3* specifically within atrial and ventricular endocardial cell populations.

**Figure 1–figure supplement 2.**
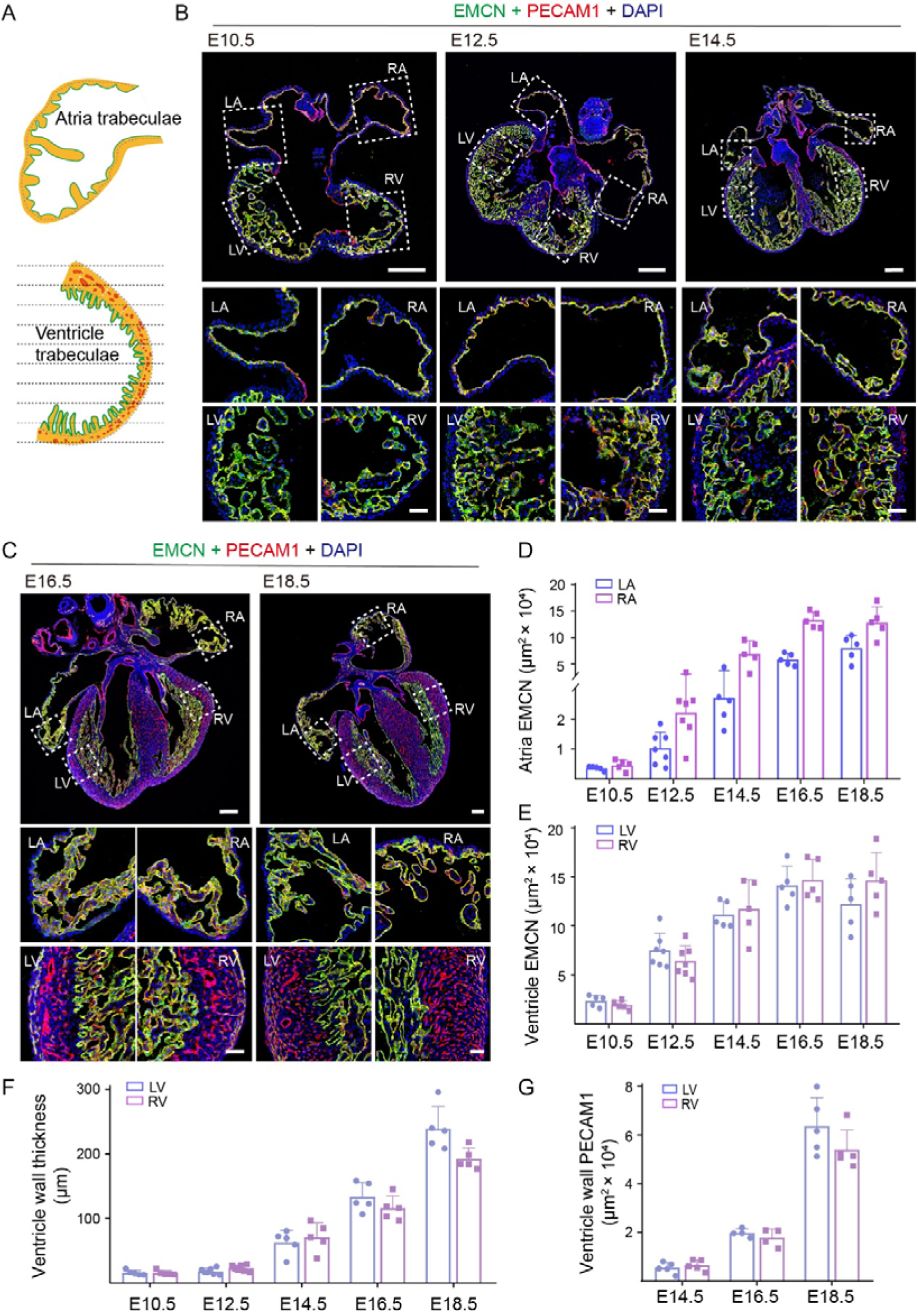
Analysis of atrial and ventricular chamber morphogenesis during the murine embryonic development. **A.** Schematic illustration of quantitative analysis for cardiac wall. Ten horizontal lines were evenly laid on the images, and the thickness of ventricular wall (epicardial side) crossed with these lines were measured and analyzed using Image Pro Plus (Media Cybernetics, Inc., Bethesda, MD). **B-C**. Analysis of murine atrial and ventricular trabeculae by immunostaining for EMCN (green), PECAM1 (red) and DAPI (blue) from E10.5-E18. Note that ventricular trabeculae were already present at E10.5, while the atrial trabeculation began around E12.5. **D-G**. Quantification of the EMCN-positive areas in the atrial and ventricular trabeculae (**D-E**), the ventricular wall thickness (**F**) and the PECAM1-positive area in ventricular walls (**G**) at the different embryonic stages. The quantitation data are shown in Supplemental Table 1. In D-G, values are represented as means ± SD of at least three independent technical replicates. Scale bar: 200 μm in B (above) and C (above); 50 μm in B (below) and C (below).

**Figure 2–figure supplement 1.**
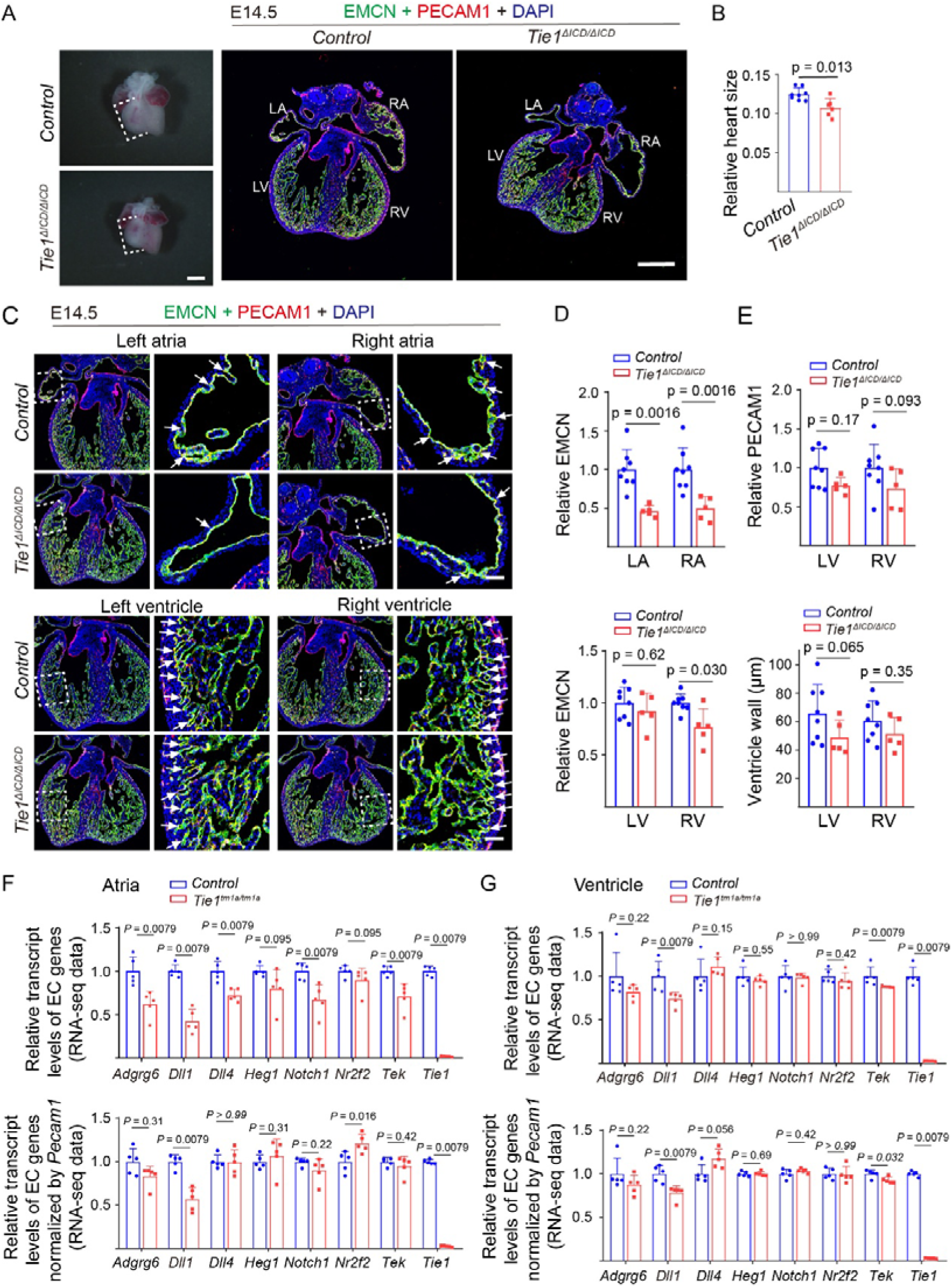
Defective atrial trabeculation after *Tie1* deletion (*Tie1*^Δ*ICD/*Δ*ICD*^). **A-C.** Analysis of cardiac trabeculae of *Tie1*^Δ*ICD/*Δ*ICD*^ and littermate controls by immunostaining for EMCN (green), PECAM1 (red) and DAPI (blue) at E14.5 **(A,C)**. The *Tie1* mutant hearts were smaller and atrial trabeculae were absent, while ventricular trabeculae were slightly sparse. Quantification of the relative heart size at E14.5 (**B**, *Tie1*^Δ*ICD/*Δ*ICD*^: 0.11±0.011, n=6; *Control*: 0.13±0.0078, n=8; *P*=0.013.). **D-E**. Quantification of the EMCN-positive area in the atrial and ventricular trabeculae (**D,** LA: *Tie1*^Δ*ICD/*Δ*ICD*^: 0.46±0.071, n=5; *Control*: 1.00±0.26, n=8; *P*=0.0016. RA: *Tie1*^Δ*ICD/*Δ*ICD*^: 0.50±0.15, n=5; *Control*: 1.00±0.28, n=8; *P*=0.0016. LV: *Tie1*^Δ*ICD/*Δ*ICD*^: 0.92±0.17, n=5; *Control*: 1.00±0.15, n=8; *P*=0.62. RV: *Tie1*^Δ*ICD/*Δ*ICD*^: 0.77±0.17, n=5; *Control*: 1.00±0.085, n=8; *P*=0.030.). Quantification of the PECAM1-positive area in ventricular walls (**E,** LV: *Tie1*^Δ*ICD/*Δ*ICD*^: 0.77±0.10, n=5; *Control*: 1.00±0.25, n=8; *P*=0.17. RV: *Tie1*^Δ*ICD/*Δ*ICD*^: 0.73±0.25, n=5; *Control*: 1.00±0.30, n=8; *P*=0.093.). Quantification of the ventricular wall thickness (**E,** LV: *Tie1*^Δ*ICD/*Δ*ICD*^: 49.00±11.97 µm, n=5; *Control*: 65.72±20.43 µm, n=8; *P*=0.065. RV: *Tie1*^Δ*ICD/*Δ*ICD*^: 51.28±11.59 µm, n=5; *Control*: 60.68±14.08 µm, n=8; *P*=0.35.) at E14.5. Except the ventricular wall thickness, the quantification data of mutant mice was normalized against that of littermate control mice. **F-G**. Alteration of the trabeculation related genes upon *Tie1* deficiency in atria (F) and ventricles (G). The quantitative results are shown in Supplemental Table 3 and Supplemental Table 4. In B, D-G statistical analysis was performed using the unpaired nonparametric Mann-Whitney U test. Values are represented as means ± SD of at least three independent technical replicates. LA: Left atria, RA: Right atria, LV: Left ventricle, RV: Right ventricle; Scale bar: 400 μm in A, 50 μm in C.

**Figure 2–figure supplement 2.**
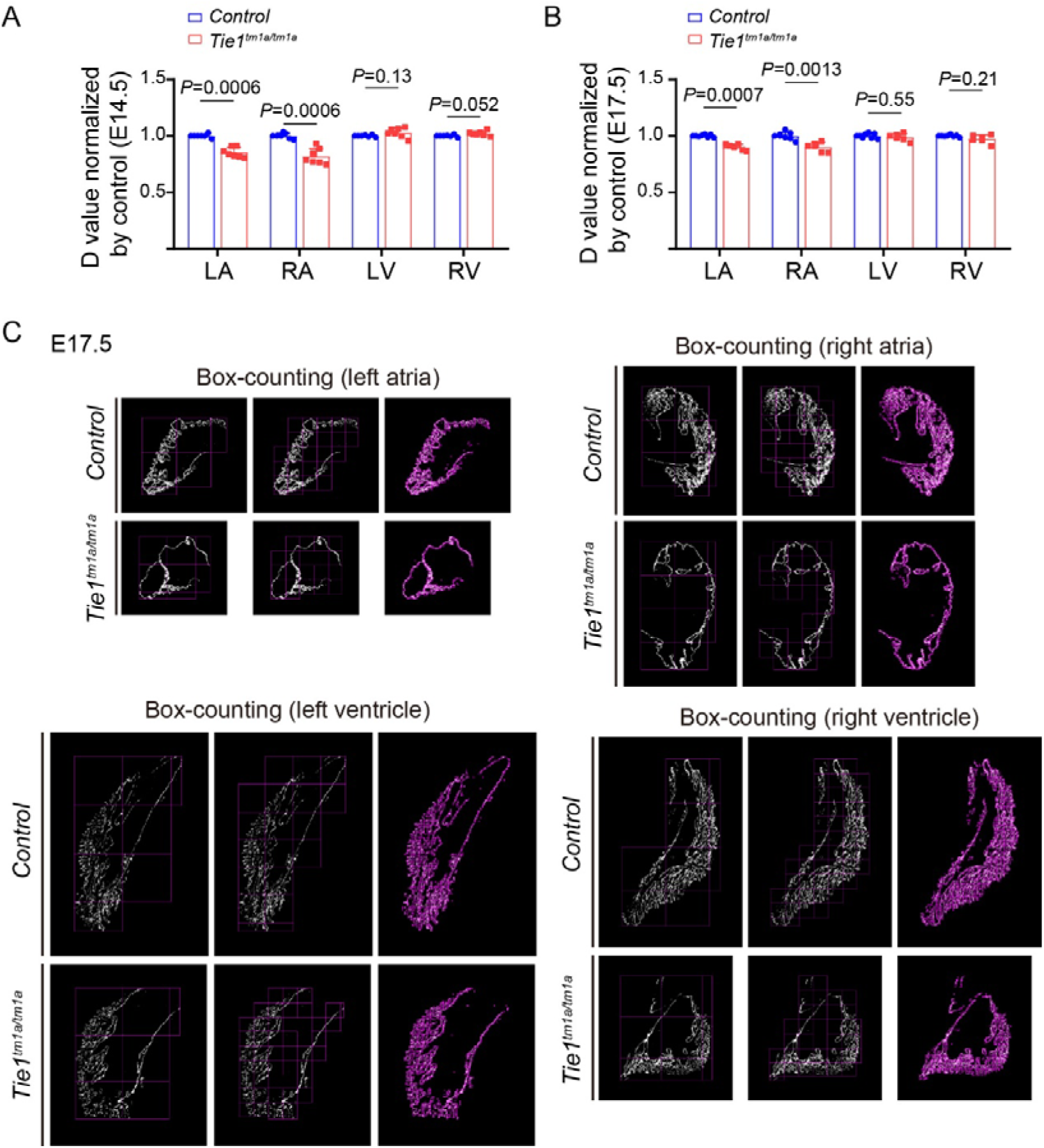
Fractal dimension analysis of atrial and ventricular trabeculae from the *Tie1* mutant and littermate control mice. A-B. Quantification of the box-counting fractal dimension of atrial and ventricular trabeculae from *Tie1^tm1a/tm1a^* and littermate controls at E14.5 (A, Supplemental Table 2) and E17.5 (B, Supplemental Table 2). C. Representative images used for the quantification of the box-counting fractal dimension of atrial and ventricular trabeculae from *Tie1* mutant mice at E17.5.

**Figure 2–figure supplement 3.**
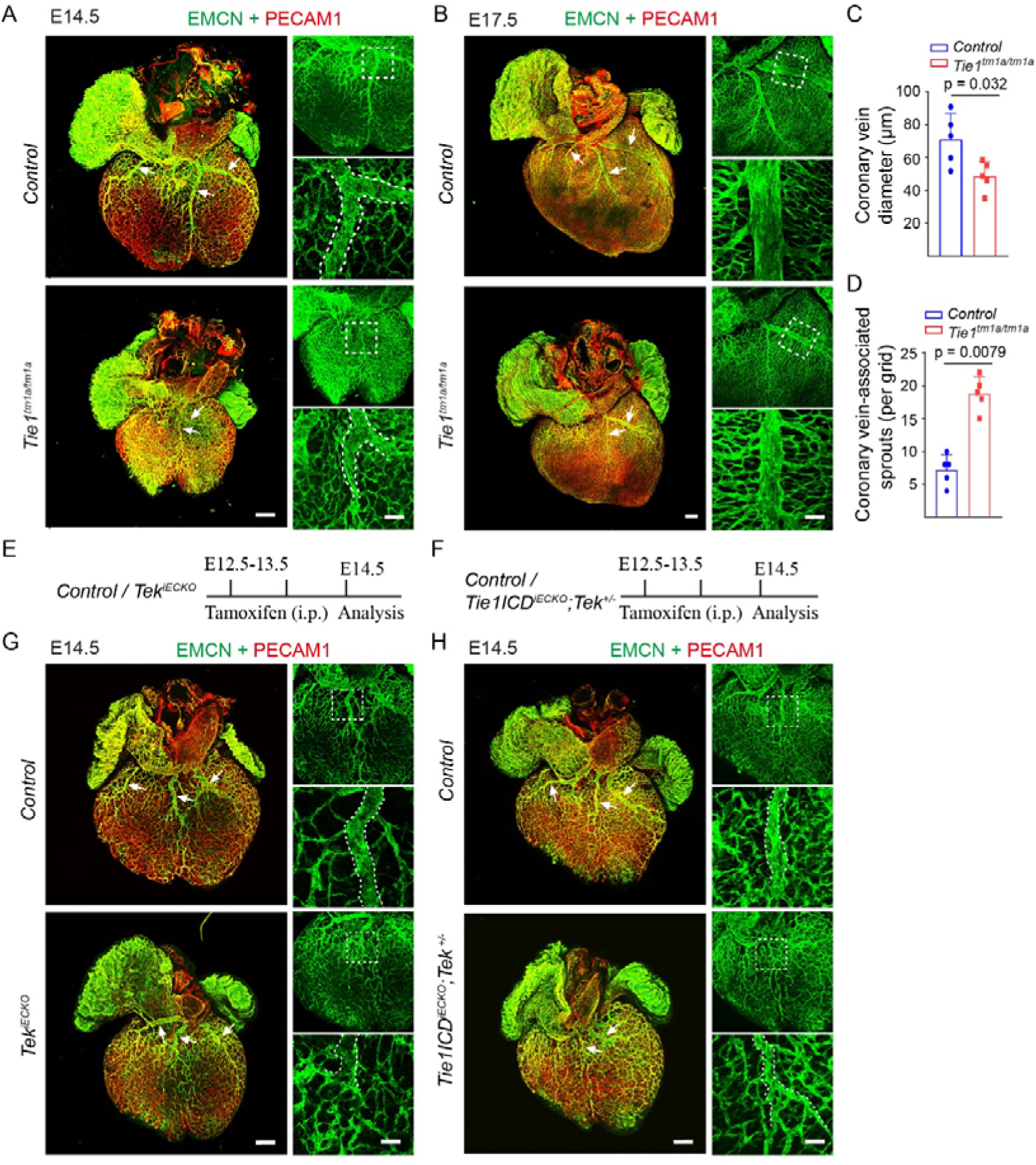
Defective coronary vein formation after TIE1/TIE2 deficiency or insufficiency. **A-B.** Visualization of coronary vein (dorsal) of *Tie1^tm1a/tm1a^*and littermate controls by whole-mount immunostaining for EMCN (green) and PECAM1 (red) at E14.5 (**A**) and E17.5 (**B**). TIE1 deficiency (*Tie1^tm1a/tm1a^*) led to a decrease in the coronary vein diameter but an increase with the vein-associated angiogenic sprouting (E17.5). **C-D.** Quantitative analysis of coronary vein diameter (**C**, *Tie1^tm1a/tm1a^*: 48.80±9.12, n=5; *Control*: 71.21±15.65, n=5; *P*=0.032.) and sprouts (**D**, *Tie1^tm1a/tm1a^*: 18.80±2.59, n=5; *Control*: 7.20±2.28, n=5; *P*=0.0079.) at E17.5. **E-H.** Analysis of coronary vein (dorsal) of *Tek^iECKO^* (**E,G**), *Tie1ICD^iECKO^;Tek^+/-^*(**F,H**) and littermate controls by whole-mount immunostaining for EMCN (green) and PECAM1 (red) at E14.5. The tamoxifen administration by intraperitoneal (i.p.) and analysis scheme were shown in **E-F**. White arrows point to the abnormal and delayed coronary vein formation in *Tie1*, *Tek* and *Tie1/Tek* mutant mice. In C and D, statistical analysis was performed using the unpaired nonparametric Mann-Whitney U test. Values are represented as means ± SD of at least three independent technical replicates. Scale bar: 200 μm in left of A, B, G, H; 50 μm in right of A, B, G, H.

**Figure 3–figure supplement 1.** 3D views of the wholemount immunostaining of hearts from the *Tie1^tm1a/tm1a^* and littermate controls as shown in Figure 3A.

**Supplement Table 1.** Quantitation of the trabecular area (EMCN^+^), blood vessel density (PECAM1^+^) in ventricular wall and ventricular wall thickness at the different stages of murine embryonic development (E10.5-E18.5). Values are represented as means ± SD of at least three independent technical replicates.

| | | E10.5<br>(n=5,<br>mean±sd) | E12.5<br>(n=7,<br>mean±sd) | E14.5<br>(n=5,<br>mean±sd) | E16.5<br>(n=5,<br>mean±sd) | E18.5<br>(n=5,<br>mean±sd) |
| --- | --- | --- | --- | --- | --- | --- |
| **Atria**<br>**EMCN**<br>**(μm<sup>2</sup> ×**<br>**10<sup>3</sup>)** | LA | 0.36±0.066 | 1.03±0.54 | 2.70±1.04 | 5.88±1.12 | 8.02±2.39 |
| | RA | 0.47±0.18 | 2.22±0.86 | 6.91±2.51 | 13.34±1.54 | 12.89±2.90 |
| **Ventricle**<br>**EMCN**<br>**(μm<sup>2</sup> ×**<br>**10<sup>3</sup>)** | LV | 2.34±0.57 | 7.49±1.72 | 11.13±1.34 | 14.14±1.95 | 12.22±2.58 |
| | RV | 1.91±0.44 | 6.39±1.57 | 11.74±2.93 | 14.67±2.09 | 14.62±2.86 |
| **Ventricle**<br>**wall (μm)** | LV | 15.43±3.88 | 17.72±4.68 | 62.51±18.57 | 132.76±22.15 | 238.81±34.11 |
| | RV | 15.21±3.68 | 22.64±4.95 | 70.65±22.17 | 115.72±18.71 | 191.94±16.55 |
| **Ventricle**<br>**wall**<br>**PECAM1**<br>**(μm<sup>2</sup> ×**<br>**10<sup>3</sup>)** | LV | - | - | 0.51±0.23 | 1.95±0.17<br>(n=4) | 6.34±1.17 |
| | RV | - | - | 0.62±0.23 | 1.75±0.36<br>(n=4) | 5.36±0.82 |

**Supplement Table 2.** Quantification of the fractal dimension of trabecular in *Tie1^tm1a/tm1a^*and littermate controls by using box-counting analysis at E14.5 and E17.5. Statistical analysis was performed using the unpaired nonparametric Mann- Whitney U test. Values are represented as means ± SD of at least three technical replicates.

|  |  | LA | RA | LV | RV |
| --- | --- | --- | --- | --- | --- |
| **Relative D**<br>**value**<br>**(E14.5)** | *Control*<br>(n=7, mean±sd) | 1.00±0.016 | 1.00±0.024 | 1.00±0.0082 | 1.00±0.0090 |
| | *Tie1*<sup>tm1a/tm1a</sup><br>(n=7, mean±sd) | 0.85±0.045 | 0.82±0.070 | 1.03±0.042 | 1.02±0.021 |
|  | *P* | 0.00060 | 0.00060 | 0.13 | 0.052 |
| **Relative D**<br>**value** | *Control*<br>(n=8, mean±sd) | 1.00±0.011 | 1.00±0.032 | 1.00±0.020 | 1.00±0.0087 |
| (E17.5) | *Tie1*<sup>tm1a/tm1a</sup><br>(n=6, mean±sd) | 0.90±0.026 | 0.90±0.041 | 0.99±0.032 | 0.97±0.037 |
|  | *P* | 0.00070 | 0.0013 | 0.55 | 0.21 |

**Supplement Table 3.** Alteration of the transcript levels of atrial trabeculation-related genes after *Tie1* deficiency (*Tie1^tm1a/tm1a^* and control mice, E14.5). Statistical analysis was performed using the unpaired nonparametric Mann-Whitney U test. Values are represented as means ± SD of at least three technical replicates.

|  | Relative transcript levels of EC genes |  |  | Relative transcript levels of EC genes normalized by *Pecam1* |  |  |
| --- | --- | --- | --- | --- | --- | --- |
|  | *Control*<br>(n = 5,<br>mean±sd) | *Tie1*<sup>tm1a/tm1a</sup><br>(n = 5,<br>mean±sd) | *P* | *Control*<br>(n = 5,<br>mean±sd) | *Tie1*<sup>tm1a/tm1a</sup><br>(n = 5,<br>mean±sd) | *P* |
| *Adgrg6* | 1.00±0.16 | 0.62±0.15 | 0.0079 | 1.00±0.15 | 0.83±0.12 | 0.31 |
| *Dll1* | 1.00±0.075 | 0.42±0.13 | 0.0079 | 1.00±0.079 | 0.57±0.13 | 0.0079 |
| *Dll4* | 1.00±0.11 | 0.72±0.069 | 0.0079 | 1.00±0.078 | 1.00±0.14 | >0.99 |
| *Heg1* | 1.00±0.064 | 0.80±0.22 | 0.095 | 1.00±0.079 | 1.07±0.19 | 0.31 |
| *Notch1* | 1.00±0.097 | 0.67±0.17 | 0.0079 | 1.00±0.044 | 0.90±0.13 | 0.22 |
| *Nr2f2* | 1.00±0.066 | 0.90±0.14 | 0.095 | 1.00±0.12 | 1.21±0.10 | 0.016 |
| *Tek* | 1.00±0.056 | 0.71±0.15 | 0.0079 | 1.00±0.048 | 0.95±0.11 | 0.42 |
| *Tie1* | 1.00±0.059 | 0.020±0.0077 | 0.0079 | 1.00±0.026 | 0.026±0.0036 | 0.0079 |

**Supplement Table 4.** Alteration of the transcript levels of ventricular trabeculation- related genes after *Tie1* deficiency (*Tie1^tm1a/tm1a^* and control mice, E14.5). Statistical analysis was performed using the unpaired nonparametric Mann-Whitney U test. Values are represented as means ± SD of at least three technical replicates.

|  | Relative transcript levels of EC genes |  |  | Relative transcript levels of EC genes normalized by *Pecam1* |  |  |
| --- | --- | --- | --- | --- | --- | --- |
|  | *Control*<br>(n = 5,<br>mean±sd) | *Tie1*<sup>tm1a/tm1a</sup><br>(n = 5,<br>mean±sd) | *P* | *Control*<br>(n = 5,<br>mean±sd) | *Tie1*<sup>tm1a/tm1a</sup><br>(n = 5,<br>mean±sd) | *P* |
| *Adgrg6* | 1.00±0.28 | 0.82±0.081 | 0.22 | 1.00±0.18 | 0.87±0.11 | 0.22 |
| *Dll1* | 1.00±0.17 | 0.74±0.079 | 0.0079 | 1.00±0.093 | 0.78±0.080 | 0.0079 |
| *Dll4* | 1.00±0.20 | 1.11±0.11 | 0.15 | 1.00±0.10 | 1.18±0.11 | 0.056 |
| *Heg1* | 1.00±0.11 | 0.95±0.050 | 0.55 | 1.00±0.028 | 1.00±0.026 | 0.69 |
| *Notch1* | 1.00±0.13 | 0.98±0.054 | >0.99 | 1.00±0.053 | 1.04±0.026 | 0.42 |
| *Nr2f2* | 1.00±0.080 | 0.95±0.096 | 0.42 | 1.00±0.065 | 0.99±0.094 | >0.99 |
| *Tek* | 1.00±0.11 | 0.88±0.010 | 0.0079 | 1.00±0.044 | 0.93±0.037 | 0.032 |
| *Tie1* | 1.00±0.11 | 0.027±0.0024 | 0.0079 | 1.00±0.030 | 0.028±0.0031 | 0.0079 |

**Supplement Table 5.** The differential transcript levels of cardiac trabeculation-related genes between atria and ventricles (E14.5, based on the RNA-seq analysis of atria and ventricles separately). Statistical analysis was performed using the unpaired nonparametric Mann-Whitney U test. Values are represented as means ± SD of at least three technical replicates.

|  | Relative transcript levels of EC genes |  |  | Relative transcript levels of EC genes normalized by *Pecam1* |  |  |
| --- | --- | --- | --- | --- | --- | --- |
| | Atria<br>(n = 5,<br>mean±sd) | Ventricle<br>(n = 5,<br>mean±sd) | *P* | Atria<br>(n = 5,<br>mean±sd) | Ventricle<br>(n = 5,<br>mean±sd) | *P* |
| *Adgrg6* | 1.00±0.16 | 0.38±0.10 | 0.0079 | 1.00±0.15 | 0.37±0.067 | 0.0079 |
| *Dll1* | 1.00±0.075 | 0.68±0.12 | 0.0079 | 1.00±0.079 | 0.66±0.062 | 0.0079 |
| *Dll4* | 1.00±0.11 | 1.30±0.26 | 0.016 | 1.00±0.078 | 1.27±0.13 | 0.0079 |
| *Heg1* | 1.00±0.064 | 0.94±0.10 | 0.15 | 1.00±0.079 | 0.92±0.026 | 0.032 |
| *Notch1* | 1.00±0.097 | 0.93±0.12 | 0.84 | 1.00±0.044 | 0.92±0.048 | 0.032 |
| *Nr2f2* | 1.00±0.066 | 0.25±0.020 | 0.0079 | 1.00±0.12 | 0.24±0.016 | 0.0079 |
| *Tek* | 1.00±0.056 | 0.56±0.062 | 0.0079 | 1.00±0.048 | 0.55±0.024 | 0.0079 |
| *Tie1* | 1.00±0.059 | 0.73±0.079 | 0.0079 | 1.00±0.026 | 0.72±0.021 | 0.0079 |

## References

1. Tian, Y. and E.E. Morrisey, Importance of myocyte-nonmyocyte interactions in cardiac development and disease. Circ Res, 2012. 110(7): p. 1023–34.

2. Meilhac, S.M. and M.E. Buckingham, The deployment of cell lineages that form the mammalian heart. Nat Rev Cardiol, 2018. 15(11): p. 705–724.

3. Moorman, A.F. and V.M. Christoffels, Cardiac chamber formation: development, genes, and evolution. Physiol Rev, 2003. 83(4): p. 1223–67.

4. Tran, Y.T.H., D. Saha, and G. Del Monte-Nieto, Cardiac trabeculation in vertebrates: Convergent evolution or evolutionary adaptations associated with heart complexity? Semin Cell Dev Biol, 2025. 172: p. 103622.

5. Haack, T. and S. Abdelilah-Seyfried, The force within: endocardial development, mechanotransduction and signalling during cardiac morphogenesis. Development, 2016. 143(3): p. 373–86.

6. Rhee, S., et al., Endocardial/endothelial angiocrines regulate cardiomyocyte development and maturation and induce features of ventricular non-compaction. Eur Heart J, 2021. 42(41): p. 4264–4276.

7. Captur, G., et al., Formation and Malformation of Cardiac Trabeculae: Biological Basis, Clinical Significance, and Special Yield of Magnetic Resonance Imaging in Assessment. Can J Cardiol, 2015. 31(11): p. 1325–37.

8. Gunawan, F., R. Priya, and D.Y.R. Stainier, Sculpting the heart: Cellular mechanisms shaping valves and trabeculae. Curr Opin Cell Biol, 2021. 73: p. 26–34.

9. Samsa, L.A., B. Yang, and J. Liu, Embryonic cardiac chamber maturation: Trabeculation, conduction, and cardiomyocyte proliferation. Am J Med Genet C Semin Med Genet, 2013. 163C(3): p. 157–68.

10. Wu, B., et al., Endocardial cells form the coronary arteries by angiogenesis through myocardial-endocardial VEGF signaling. Cell, 2012. 151(5): p. 1083–96.

11. Lai, D., et al., Neuregulin 1 sustains the gene regulatory network in both trabecular and nontrabecular myocardium. Circ Res, 2010. 107(6): p. 715–27.

12. Grego-Bessa, J., et al., Notch signaling is essential for ventricular chamber development. Dev Cell, 2007. 12(3): p. 415–29.

13. Watanabe, Y., et al., Activation of Notch1 signaling in cardiogenic mesoderm induces abnormal heart morphogenesis in mouse. Development, 2006. 133(9): p. 1625–34.

14. Del Monte-Nieto, G., et al., Control of cardiac jelly dynamics by NOTCH1 and NRG1 defines the building plan for trabeculation. Nature, 2018. 557(7705): p. 439–445.

15. Miao, L., et al., Tunneling nanotube-like structures regulate distant cellular interactions during heart formation. Science, 2025. 387(6739): p. eadd3417.

16. Lockhart, M., et al., Extracellular matrix and heart development. Birth Defects Res A Clin Mol Teratol, 2011. 91(6): p. 535–50.

17. Kern, C.B., et al., Reduced versican cleavage due to Adamts9 haploinsufficiency is associated with cardiac and aortic anomalies. Matrix Biol, 2010. 29(4): p. 304–16.

18. Kim, K.H., et al., Myocardial Angiopoietin-1 Controls Atrial Chamber Morphogenesis by Spatiotemporal Degradation of Cardiac Jelly. Cell Rep, 2018. 23(8): p. 2455–2466.

19. Augustin, H.G., et al., Control of vascular morphogenesis and homeostasis through the angiopoietin-Tie system. Nat Rev Mol Cell Biol, 2009. 10(3): p. 165–77.

20. Saharinen, P., L. Eklund, and K. Alitalo, Therapeutic targeting of the angiopoietin-TIE pathway. Nat Rev Drug Discov, 2017. 16(9): p. 635–661.

21. Qu, X., C. Harmelink, and H.S. Baldwin, Tie2 regulates endocardial sprouting and myocardial trabeculation. JCI Insight, 2019. 5.

22. Leppanen, V.M., P. Saharinen, and K. Alitalo, Structural basis of Tie2 activation and Tie2/Tie1 heterodimerization. Proc Natl Acad Sci U S A, 2017. 114(17): p. 4376–4381.

23. Puri, M.C., et al., The receptor tyrosine kinase TIE is required for integrity and survival of vascular endothelial cells. Embo J, 1995. 14(23): p. 5884–91.

24. Qu, X., et al., Abnormal embryonic lymphatic vessel development in Tie1 hypomorphic mice. Development, 2010. 137(8): p. 1285–95.

25. Cao, X., et al., Endothelial TIE1 Restricts Angiogenic Sprouting to Coordinate Vein Assembly in Synergy With Its Homologue TIE2. Arterioscler Thromb Vasc Biol, 2023. 43(8): p. e323–e338.

26. Qu, X., et al., Loss of flow responsive Tie1 results in ImpairedAortic valve remodeling. Dev Biol, 2019.

27. Shen, B., et al., Genetic dissection of tie pathway in mouse lymphatic maturation and valve development. Arterioscler Thromb Vasc Biol, 2014. 34(6): p. 1221–30.

28. Chu, M., et al., Angiopoietin receptor Tie2 is required for vein specification and maintenance via regulating COUP-TFII. Elife, 2016. 5:e21032.

29. Pereira, F.A., et al., The orphan nuclear receptor COUP-TFII is required for angiogenesis and heart development. Genes Dev, 1999. 13(8): p. 1037–49.

30. You, L.R., et al., Suppression of Notch signalling by the COUP-TFII transcription factor regulates vein identity. Nature, 2005. 435(7038): p. 98–104.

31. Wu, S.P., et al., Atrial identity is determined by a COUP-TFII regulatory network. Dev Cell, 2013. 25(4): p. 417–26.

32. Michelini, S., et al., TIE1 as a Candidate Gene for Lymphatic Malformations with or without Lymphedema. Int J Mol Sci, 2020. 21(18).

33. Brouillard, P., et al., Loss-of-function mutations of the TIE1 receptor tyrosine kinase cause late-onset primary lymphedema. J Clin Invest, 2024. 134(14).

34. Feng, W., et al., Single-cell transcriptomic analysis identifies murine heart molecular features at embryonic and neonatal stages. Nat Commun, 2022. 13(1): p. 7960.

35. Okabe, K., et al., Neurons limit angiogenesis by titrating VEGF in retina. Cell, 2014. 159(3): p. 584–96.

36. Cao, X., et al., A Genetically Engineered Mouse Model of Venous Anomaly and Retinal Angioma-like Vascular Malformation. Bio Protoc, 2021. 11(15): p. e4117.

37. Kalucka, J., et al., Single-Cell Transcriptome Atlas of Murine Endothelial Cells. Cell, 2020. 180(4): p. 764–779 e20.

38. Liberzon, A., et al., The Molecular Signatures Database (MSigDB) hallmark gene set collection. Cell Syst, 2015. 1(6): p. 417–425.

39. Fatkin, D., I.G. Huttner, and R. Johnson, Genetics of atrial cardiomyopathy. Curr Opin Cardiol, 2019. 34(3): p. 275–281.

40. Arita, Y., et al., Myocardium-derived angiopoietin-1 is essential for coronary vein formation in the developing heart. Nat Commun, 2014. 5: p. 4552.

41. Sato, T.N., et al., Distinct roles of the receptor tyrosine kinases Tie-1 and Tie-2 in blood vessel formation. Nature, 1995. 376(6535): p. 70–4.

42. Saharinen, P., et al., Angiopoietins assemble distinct Tie2 signalling complexes in endothelial cell-cell and cell-matrix contacts. Nat Cell Biol, 2008. 10(5): p. 527–37.

43. Fukuhara, S., et al., Differential function of Tie2 at cell-cell contacts and cell-substratum contacts regulated by angiopoietin-1. Nat Cell Biol, 2008. 10(5): p. 513–26.

44. Albu, M., et al., Distinct mechanisms regulate ventricular and atrial chamber wall formation. Nat Commun, 2024. 15(1): p. 8159.

